# Structural architecture of amyloid-β oligomers, curvilinear protofibrils and annular assemblies, imaged by cryo-EM and cryo-ET

**DOI:** 10.1101/2024.03.01.582902

**Authors:** Ruina Liang, Andrea P. Torres-Flores, Shang Qi, Anum Khursheed, Yao Tian, Piotr Szwedziak, Mark D. Baker, Vladimir A. Volkov, Vidya C. Darbari, John H. Viles

## Abstract

Small prefibrillar structures of the amyloid-β peptide (Aβ) are believed to be central to cytotoxicity in Alzheimer’s disease, the most common form of dementia. A snapshot of these prefibrillar assemblies has therefore been characterized using a combination of cryo-ET and cryo-EM single particle analysis. This has facilitated an understanding of the relationship between, oligomers, curvilinear protofibrils and annular assemblies. A highly consistent diameter for all curvilinear protofibrils and oligomers of 28 Å, indicates that these assemblies are simply structural extensions from the smaller oligomers. Furthermore, their basic crosssection suggests mature amyloid fibrils might be initiated by the lateral binding of two curvilinear protofibrils. *Ab-initio* 3D reconstruction also reveals ring-shaped annular assemblies. These possess a central internal channel, *ca*. 14 Å in diameter and 54 Å long, which is capable of traversing lipid membranes. Large conductance recorded using patch-clamp electrophysiology, match the internal diameter of the Aβ annular architecture.

## INTRODUCTION

The amyloid cascade hypothesis describes the self-assembly of a 42 amino-acid peptide, Amyloid-β (Aβ_42_), and accounts for the most common form of dementia; Alzheimer’s disease (AD)(1, 2). Recent success in clinical trials targeted at Aβ clearance supports the amyloid cascade hypothesis(3). Thus, understanding the self-assembly of monomeric Aβ through to: oligomeric; curvilinear-protofibrillar; annular and fibrillar structures is a key question for AD and other protein misfolding diseases(4-7). The prefibrillar assemblies are generally believed to be the cytotoxic form of Aβ_42_ and to be responsible for the cascade of events which leads to dementia(4, 5, 8-12). In particular, prefibrillar Aβ_42_ has been shown to disrupt membrane integrity which causes unregulated cellular Ca^2+^ influx(13-16). While in turn the anionic phospholipid bilayer has been shown to accelerate Aβ fibril formation(17).

Aβ amyloid fibrils with a repeating intramolecular cross-β motif have been the subject of numerous structural studies at near atomic-resolution(18-24). However, these fibril structures are not thought to be cytotoxic, although they are able to surface-catalyse the formation of toxic oligomers and curvilinear protofibrils(15, 25). Obtaining high resolution structures of the smaller, metastable, prefibrillar assemblies, is more of a challenge(26, 27), biophysical approaches have included NMR(6, 28-30) and AFM(31-34). Oligomers and curvilinear protofibrils, rather than mature fibrils, insert and carpet lipid membranes(13, 14, 35), this is believed to induce membrane permeability(15, 16). Annular ring structures composed of Aβ have been suggested to span lipid bilayers and form ion-channel pores that cause cellular Ca^2+^ influx(36-39). However, these annular pore-like structures remain controversial, as it is not clear, from the AFM(37, 40) and heavy metal negatively-stained 2D TEM images(41), whether the indentation imaged in these assemblies can traverse the membrane to form a channel-pore.

Here we have used a combination of cryo-EM single particle analysis and cryo-ET to obtain 3D images of Aβ assemblies at the nanoscale, under near native conditions. We present evidence to indicate oligomers and curvilinear protofibrils are a continuum of the same structure, and we argue that the lateral self-association of two curvilinear protofibrils forms a basic stable mature amyloid fibril. We reveal the molecular architecture of the ring-like annular assemblies in near native conditions, to show how the internal channels relate to Aβ ion-channel pore conductance. The molecular architecture of prefibrillar assemblies has revealed the relationship between different meta-stable structures and membrane permeability.

## RESULTS AND DISCUSSION

Cryo-EM single particle analysis has been used to characterise prefibrillar Aβ_42_ assemblies, 14000 movies were collected on the 300kV Titan Krios at a pixel size of 1.1 Å/pixel. The Aβ_42_ preparations have been studied at the end of the lag-phase before appreciable fibril formation. After screening to establish appropriate conditions, using a range of Aβ_42_ concentrations and incubation times, a 300 µM and 30 min incubation time from monomer was determined to be appropriate. These metastable structures rapidly form fibrils so there was no attempt to isolate or separate prefibrillar assemblies any further, instead, we studied these assemblies as a heterogeneous mixture. In this way we have obtained a ‘snapshot’ of the assembly process containing all the assembly forms, from appreciable monomers through to mature amyloid fibrils. Approximately three million particles were picked using non-templated ‘cryolo’ auto-picking, a typical micrograph is shown in Figure S1. Following the removal of fibrils and artifacts, 2.5 million particles remained. A range of prefibrillar assemblies were apparent from rounds of 2D classification, these are summarized in Figure 1A. These include small circular oligomers 25-28 Å in diameter, through to extended curvilinear protofibrils, typically up to 140 Å in length. There are also some larger more circular structures often 65 Å in diameter. A limited number of more ordered mature fibrils, with a 60 Å diameter, were also present, Figure 1A.

**Figure 1:**
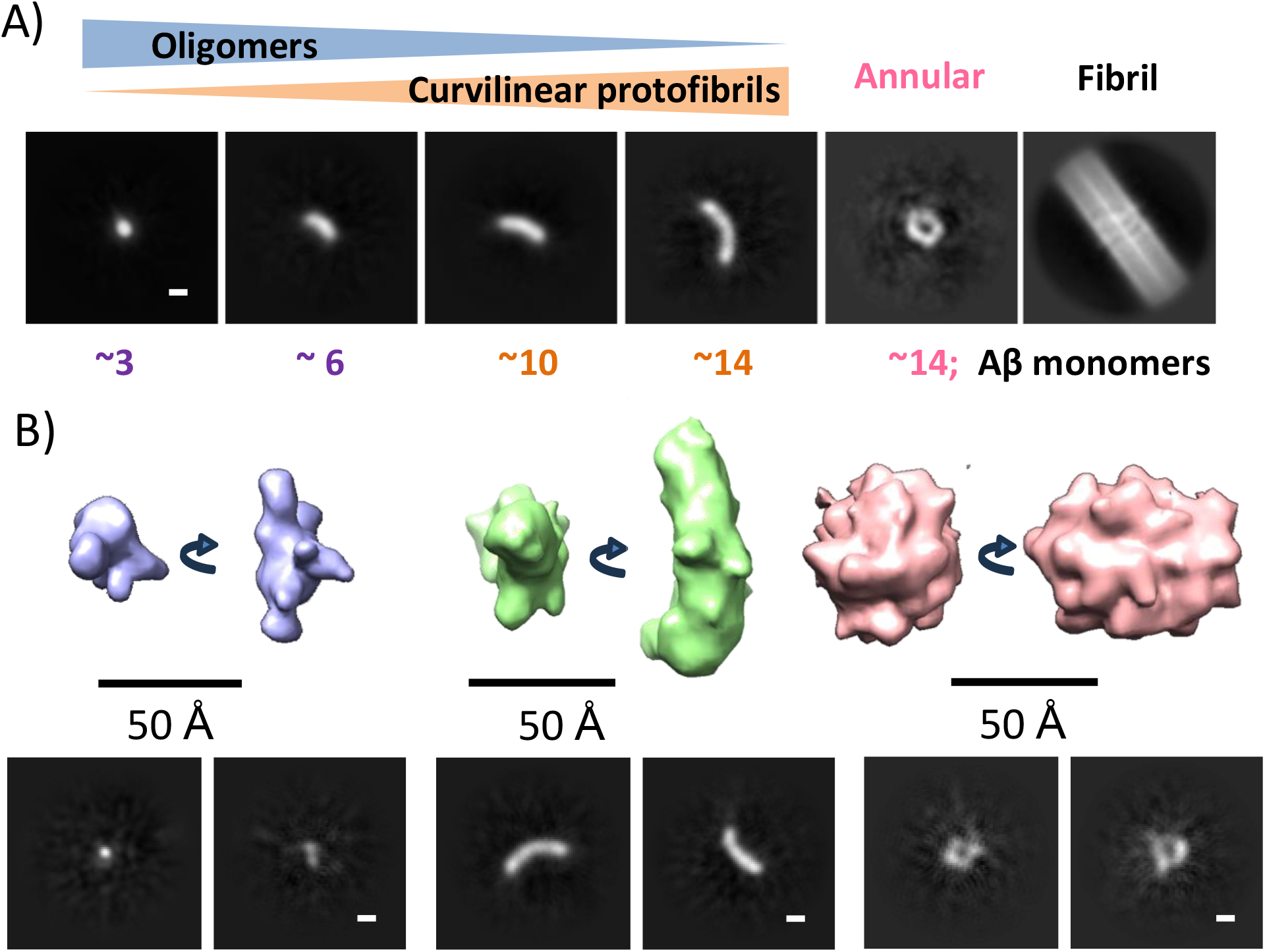
Overview of prefibrillar assemblies as a heterogeneous snap-shot, imaged by cryo-EM. Typical classification of assemblies, and approximate number of Aβ_42_ monomers is indicated (A). *Abinitio* 3D reconstruction from 2.5 million particles, 3 structures from 10 are shown. Oligomer (blue) from 180k particles; Curvilinear Protofibril (green) 322k particles and Annular assembly (pink) 298 particles. Scale Bar 20 Å for 2D class averages; 50 Å for 3D structures.

Using CryoSPARC, we performed *ab-initio* reconstruction with 2.5 million particles to generate ten heterogeneous 3D structures. These represent various intermediates in the assembly process. The range of assemblies were sufficiently different to produce distinct molecular architectures. Three of the ten structures are shown in Figure 1B: an example of an oligomer; a curvilinear protofibril; and a potential annular structure. Figure S2 summarizes the appearance of the 10 structures, the number of particles and typical 2D class averages.

### Curvilinear protofibrils imaged by cryo-EM and cryo-ET

*Cryo-Electron Microscopy:* Inspection of the ten *ab-initio* structures, Figure S2, indicated as many as seven of these structures appear as curvilinear protofibrils, with similar widths *ca*. 28 Å, but of varying length and curvature. These structures are abundant and make up 80% of the 2.5 million particles in the micrographs. A plot of the length *versus* number of curvilinear protofibrils is shown in Figure 2B, at this point in the assembly process, 95% of the particles in the class averages are less than 90 Å in length. On average the actual length of these assemblies is underestimated a little, because of the range of possible orientations for each particle. Representative 2D class averages are shown in Figure 2A. A density plot, orthogonal to the long axis, is also shown for each 2D class average, Figure 2C. These plots indicate remarkably consistent diameters 28 +/-1 Å (stand. dev.), Figure 2D. Some of the smaller globular oligomers shown on the top row, have diameters; 25 Å by 28 Å in the longer axis. The very consistent diameters, for the oligomers and curvilinear protofibrils, suggests that they are derived from the same basic fold, with increasing templated extension due to Aβ-monomer addition. We believe this is an important observation, suggesting that assemblies typically described as oligomers (top row) or curvilinear protofibrils (remaining rows) are simply the same basic structures with increasing linear addition of Aβ molecules. There are few protofibrils that exceed 200 Å in length, imaged by cryo-EM and cryo-ET, we suggest this is because longer protofibrils are prone to mechanical shearing during rotational diffusion.

**Figure 2:**
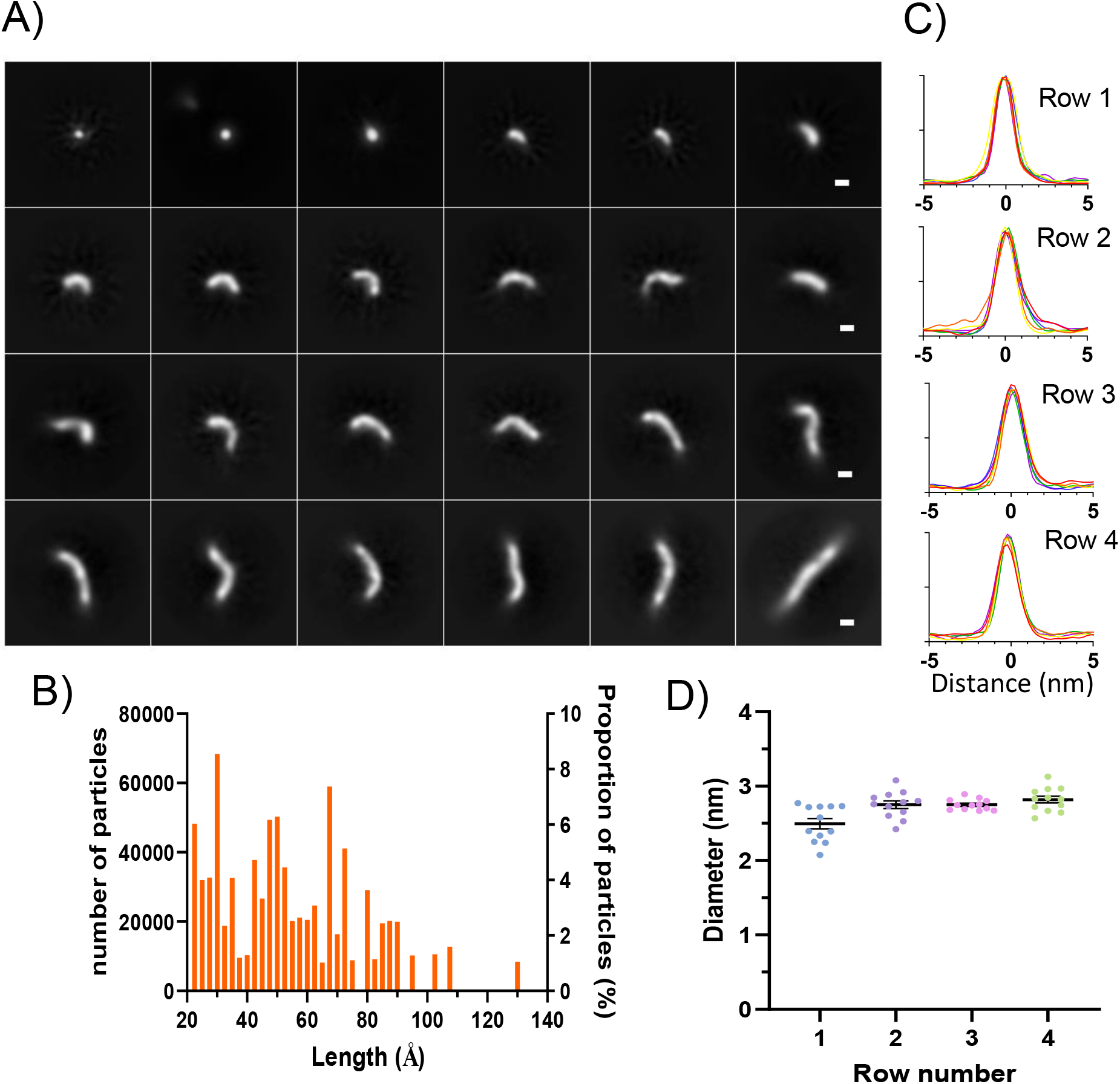
Cryo-EM 2D class averages of typical oligomers and curvilinear protofibrils. 209,000 particles in total, scale bar 20 Å. The class averages have been arranged in increasing length. Top row 25-50 Å, second row 60-80 Å; 3^rd^ row 90-105 Å; and bottom row 100-140 Å, (A). Distribution of oligomer and curvilinear protofibrils length. Taken from half a million particles; 95 % of particles are less than 90 Å long (B). Density profiles of the protofibril diameter. Six profiles for each row (Red, orange yellow left side images. Green, blue and purple on the right side) (C). Diameters observed for each row. The mean diameter for class averages in rows 2, 3 and 4 are consistently 28 +/-1 Å (standard dev.) (D).

#### Cryo-Electron Tomography

The curvilinear protofibrils are extremely variable in their curvature, indicating that generating 3D structures from single particle averaging is problematic, particularly as the structures become extended. These types of structures lend themselves to complementary cryo-ET, to obtain images of single particles. 3D tomograms of prefibrillar structures are shown in Figure 3. The 3D structures imaged in the tomograms show a very close resemblance to those determined by single particle averaging, shown in Figure 2. The cryo-ET data confirms the structures have a very consistent cross-section, with a diameter of *ca*. 28 Å. The range of lengths are mostly less than 100 Å, see supplemental figure S3A for a distribution of lengths, which is in accordance with the single particle cryo-EM data. A limited number of 2D class averages suggests branching of the curvilinear protofibrils can occur, but this is not highlighted by the 2D averaging process. Branching and irregular curves and bends in the curvilinear protofibrils are more apparent in the cryo-ET tomograms, examples of which are shown in Figure S3B.

**Figure 3:**
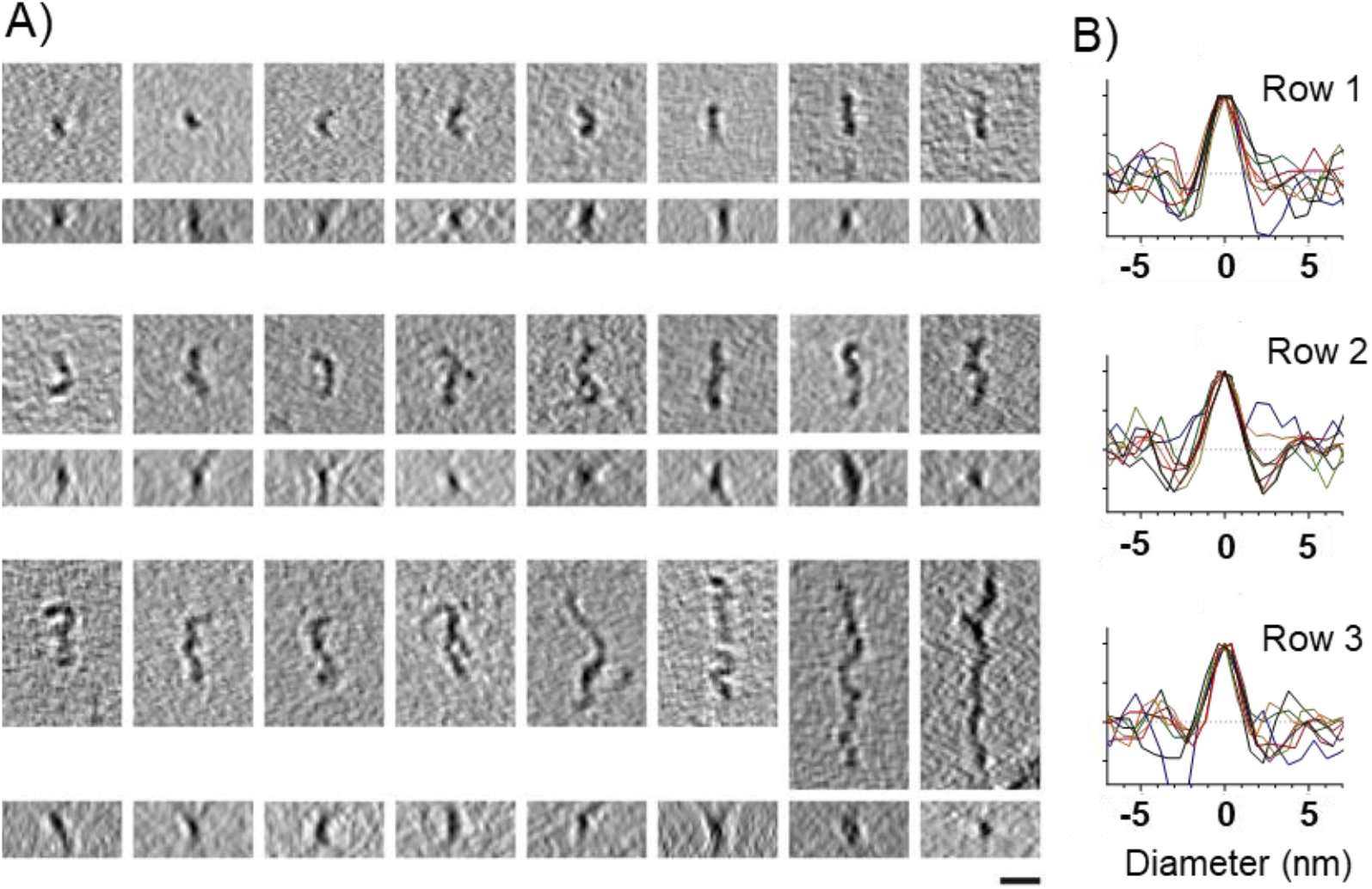
Cryo-ET 3D tomograms. (A) Typical oligomers, side and orthogonal top view are shown (top row), and curvilinear protofibrils (row 2 & 3). The cryo-ET single particles have a close resemblance to the cryo-EM class averages, shown in Figure 2. Lengths of the long axis are: Top row 25-50 Å; 2^nd^ row 60-100 Å; 3^rd^ row 100-300 Å. Images are presented as sums of 5-10 2D slices, each 7.8 Å thick. Scale Bar = 100 Å. (B) Density profiles show that the particles have a consistent diameter of *ca*. 2.8 nm.

The oligomers and curvilinear protofibrils have previously been imaged using negatively stained TEM, however the heavy-metal staining has caused their diameters to be overestimated; 6 nm is often reported(42), rather than 2.8 nm shown in Figure 2 and 3. AFM imaging has reported a height of 3 nm but has also overestimated the curvilinear protofibril widths(31, 33, 34).

*In summary*, we argue the distinction between small oligomers (28 Å in diameter) and curvilinear protofibrils is not particularly meaningful as these structures are a continuum. The curvilinear protofibrils are simply extended oligomers. Indeed, both oligomers and curvilinear protofibrils exhibit the same type of interaction with lipid membranes, both inserting into the upper-leaflet of the bilayer to the same extent and carpeting the surface causing permeability across the membrane(13-15, 43). When viewed in the round, cytotoxicity of both oligomers and curvilinear protofibrils suggest a similar membrane permeability and cytotoxicity(12, 14).

### The relationship between oligomers, curvilinear protofibrils, annular structures and fibrils

We were struck by the observation that the cross-section of the oligomers and curvilinear protofibrils, closely matches the density of half the cross-section of the basic Aβ42 amyloid fibril. The cryo-EM 3D reconstruction of a curvilinear protofibril, taken from 50k particles, is overlaid with half a basic fibril structure (pdb: 2MXU)(19) Figure 4. Numerous Aβ_42_ fibril morphologies are reported under different conditions. All contain a well conserved ‘S-shaped’ topology, between residues 15-42, Figure *4* *(18-20)*. We believe the N-terminal third, residues 1-14, are likely to be too disordered to be imaged in the curvilinear protofibrils, indeed even in the more ordered fibrils these residues are often unresolved. These residues have been shown to be dynamic and lack stable hydrogen-bonds in mature fibrils(44). We therefore postulate the curvilinear protofibrils, which are rich in β-sheet(6), form a similar S-shaped structure to fibrils, with an in-register stacking of Aβ and intermolecular hydrogen bonds. We postulate that the marked curvature in the curvilinear protofibrils is lost when fibrils are formed. The lateral association of the protofibrils adds rigidity and strength to the structure, which also facilitates extension of the fibrils with a reduction in the tendency for fragmentation. We can infer an approximate molecular weight of the assemblies shown in figure 2 and 3, assuming a crosssection slice for each Aβ molecule of 4.8 Å thickness (4.8 Å is the mid-point distance between two β-strands). The smaller oligomers of Aβ imaged are perhaps trimers-pentamers of Aβ, while a length of 48 Å suggests 10 Aβ molecules (45 kDa). A longer curvilinear protofibril of 144 Å in length, equates to 30 Aβ molecules.

**Figure 4.**
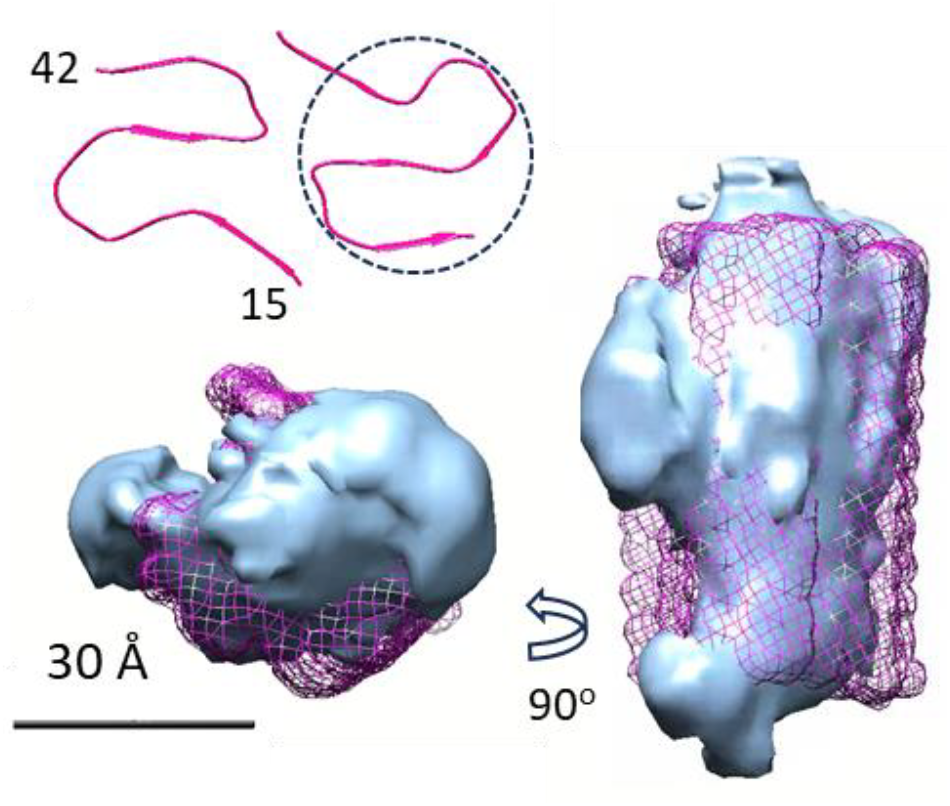
Curvilinear protofibril overlayed with fibril. Overlay of curvilinear protofibrils (cyan) with mesh for half a fibril (magenta), 3D reconstruction from 59,000 single particles. Fibril structure pdb:2MXU. Only residues 15-42 are shown in the fibril as a mesh. Residues 1-14 are often flexible in fibrils. Top left is a crosssection of the whole fibril, top-view, containing two Aβ molecules with a conserved S-shaped fold, showing β-stands; pdb:5KK3. The doted circle has a diameter of 28 Å like the curvilinear protofibrils. Scale bar: 30 Å.

We also note from single particle 2D averages that curvilinear protofibrils occasionally form tight turns that fold back on themselves to produce incomplete rings, shown in Figure S4, this is also occasionally observed in the cryo-ET tomograms. These series of incomplete ring structures suggest how annular rings, described in the next section, might also be formed from curvilinear protofibrils, while the final annular structures are 54 Å high, twice the thickness of a curvilinear protofibril. The annular structure, described in the next section, may undergo considerable rearrangement upon inserting into the hydrophobic membrane. Conceivably, the S-shape topology could stretch out while retaining its hydrogen-bonding pattern to form a β-barrel structure.

### Annular structures

The other type of prefibrillar structure is less elongated than curvilinear protofibrils, but exceeds 50 Å in all dimensions. Only two of the 10 structures generated from *ab-initio* reconstruction fit this description, less than 20% of the lag-phase particles, Figure S2.

Some of these 2D class averages (*ca*. 60 Å diameter) have a ring-like appearance, which is consistent with an annular topology, previously imaged by AFM (37, 40). Using this subset of 562k particles, three *ab-initio* structures were generated, one of these structures has a ring like appearance shown in Figure S5. The 2D class averages, representing 144k particles, is also shown in Figure S5.

This 3D reconstruction, Figure S5, has a general appearance of the annular structure suggested by AFM imaging(37, 40), with an annular ring-shape. Importantly, the 3D cryo-EM structure shown in Figure S5 contains an internal channel running through the middle of the structure. Inspection of the 2D class averages indicates that some of the classes do not match well with the general appearance of the 3D structure shown in Figure S5. To improve the resolution, the class averages that were poorly defined; too large; had density on the side of the ring; or suggested an incomplete ring, were manually removed, resulting in a reduced set of 22k particles. Inspection of the 2D class averages suggested similar structures but with more than one ring size, shown in Figure S6. Multiple ring sizes have also been suggested from the AFM images(37, 40) and have also been reported for annular structures of the amyloid protein, αsynuclein, responsible for Parkinson’s disease(45). An attempt to generate two structures *ab-initio* did not generate significantly different sizes. Thus, we took this data set and manually split them into three data sets, approximately 7 k particles each, from these three subsets, 3D structures were generated. The data set with the slightly smaller particles produced an annular structure, Figure 5. Gold standard Fourier shell correlation (FSC) indicates a resolution of 11 Å, without the use of a mask. This 3D structure generated shows a close resemblance to the set of 2D class averages from which they are derived. We have used these structures (Figure 5) and the associated 2D class averages to extract further information. In particular, the dimensions of the internal pore. The length of the pore is 54 Å, while the external diameter of the ring is between 55 and 65 Å, the approximate internal pore diameter is 10-15 Å, Figure S7. From the 3D structure a volume can be derived, which will suggest a molecular weight and Aβ stoichiometry for the annular structure. The setting of the threshold (used in Figure 5) will impact this approach but never-the-less the structure shown has a volume of *ca*. 64,000 Å^3^ which suggests a molecular weight of 53 kDa, (64,000 Å^3^ x 825 Da/Å^3^) (46) a dodecamer of Aβ_42_. The cryo-EM density might not include all the Aβ_42_ residues, for example, flexible N-terminal residues (residues 1-14) might not contribute to the cryo-structure, thus the dominant annular structure might be closer to 16 Aβ_42_molecules.

**Figure 5:**
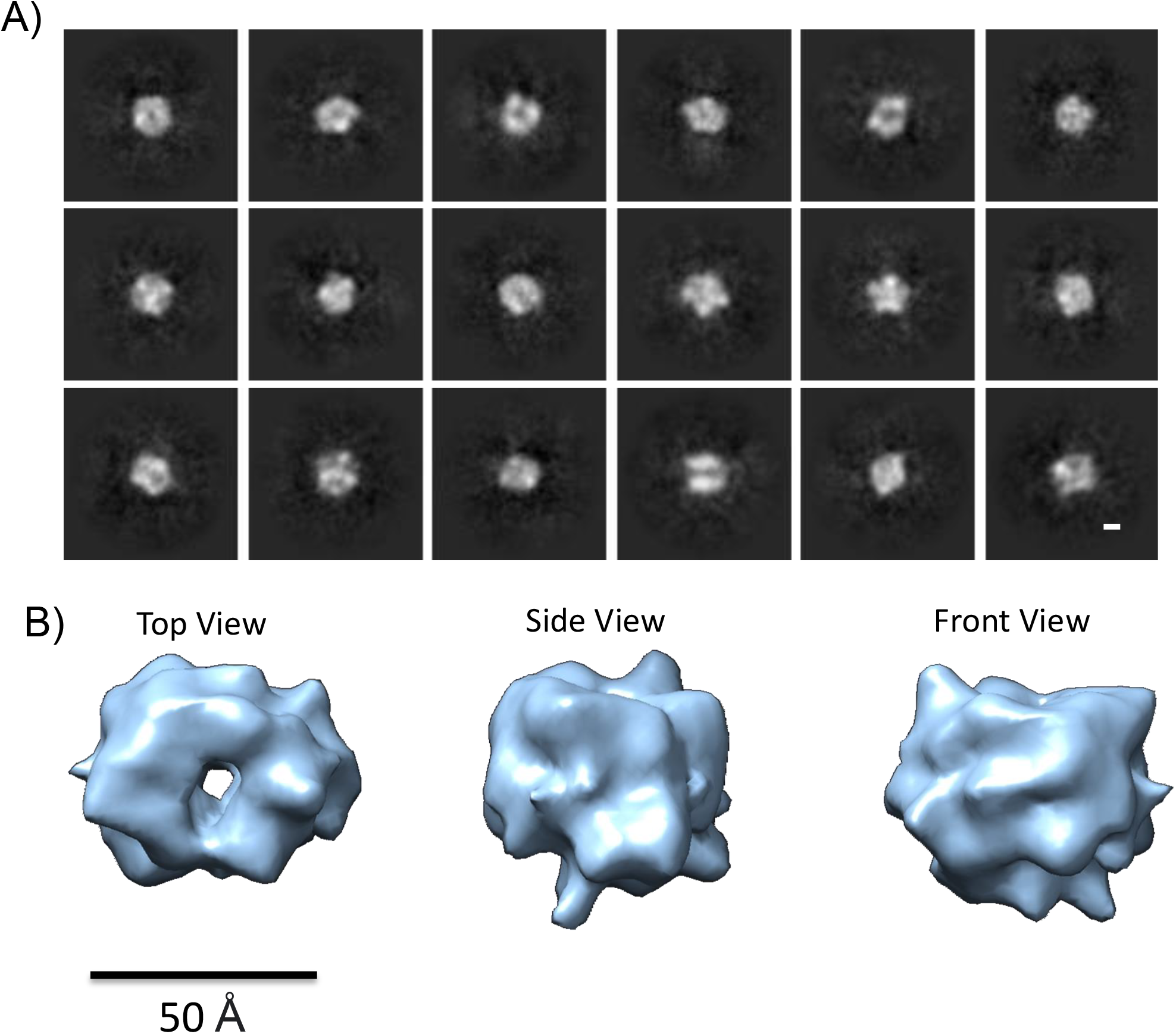
Cryo-EM 3D reconstruction of Aβ_42_ annular oligomers. 2D averages (A) and 3D reconstruction (B) from 6k single particles suggests the structure has an internal pore. External Diameter, of top view is 65-55 Å; Length of channel is 54 Å; Volume suggests 12-16 Aβ_42_ molecules. Scale bar: 50 Å for 3D structures, 20 Å for 2D averages.

High resolution structures of Aβ annular oligomers, sufficient to resolve sidechains, are a challenge because of their small size, *ca* 53 kDa, and structural heterogeneity due to the continuum of assembly pathway structures in the dataset. Furthermore, these structures may be quite dynamic and flexible with molten globule properties. Multiple channel conductance’s suggest the annular structures can transition in size, this is supported by small variations in ring-size from a survey of 2D class averages, Figure S6. The external diameter of the rings varies between 38 Å for the very smallest, while the majority (90 %) of rings are between 50-65 Å. These are the first images of annular structures under near native conditions suspended in ice. The have a strong resemblance to those imaged in a supported lipid bilayer by AFM (37, 40).

In support of the annular assemblies derived from 3D image reconstruction, we were able to identify small ring-like structures in the cryoelectron tomograms, an example of which is shown in supplemental Figure S8. The toroidal shape has an external ring diameter of *ca*. 7 nm, which closely matches the cryo-EM 3D reconstructions shown in Figure 5 and S5, the length of the internal channel appears a little smaller at *ca*. 4 nm.

Molecular dynamic simulations have predicted stable pore structures containing 12 to 20 Aβ monomers(47). The dimensions of the predicted assembly with 12 Aβ molecules(48), matches our cryo-EM structure, shown in Figure 5. A β-hairpin barrel of Aβ has been designed using a larger heptameric scaffold protein(49). The β-hairpin structures with seven Aβ molecules (PDB: 7O1Q) are compared to the annular structures determined here. Figure S9 shows the ring of the β-hairpin structure, formed with the scaffold, is smaller than for wild-type Aβ42, described in Figure 5. There has also been a much smaller β-barrel structure (3 nm external diameter) reported using a designed Aβ sequence containing cysteine mutations(50). An Aβ octamer, which forms an anti-parallel βsandwich structure imbedded in a membrane, has been described(51, 52), although the relationship to the larger annular oligomers is not clear.

### Membrane permeability, Ion-channel pore conductance and the dimensions of the annular structure

We wanted to relate the annular structure characterised here in Figure 5 to ion-channel membrane conductance, using patch-clamp electrophysiology. We and others have shown Aβ can form ion-channel pores(36-39). Indeed, we have shown only Aβ_42_ oligomers, not monomers or fibrils produce ion-channels when presented to the cellular membrane(39). Measurements here were performed using a voltage-clamped technique with an inside-out configuration, with step voltages, -80 and +80 mV. Figure 6A shows membrane conductance induced by Aβ_42_ oligomers. This was achieved by allowing a lag-phase mixture of prefibrillar assemblies to diffuse on to the extracellular surface of the excised HEK293 cellular plasma membrane. The appearance is very different from controls of buffer solution or Aβ_40_ oligomers, both of which show no membrane conductance, or occasionally small endogenous channels with conductances, 50-90 pS, Figure 6A. Aβ_42_ conductances observed are for single channels, which is indicated by conductance profiles that almost instantly become fully open or closed. The conductances produced have a particular set of properties; they are large, typically 200-500 pS. They have been shown to be ion non-selective (39) and are observed in both polarizing and depolarizing potentials. They remain open for long periods of time, often producing a ‘flickering’ appearance. The channel conductance does not usually occur immediately upon membrane exposure to Aβ_42_ oligomers, and can typically take 10 minutes for conductances to be observed after exposure. It appears the Aβ_42_ annular structures, in an aqueous environment, takes time to insert and perhaps undergo structural rearrangement within the hydrophobic lipid membrane, so as to span the bilayer.

**Figure 6:**
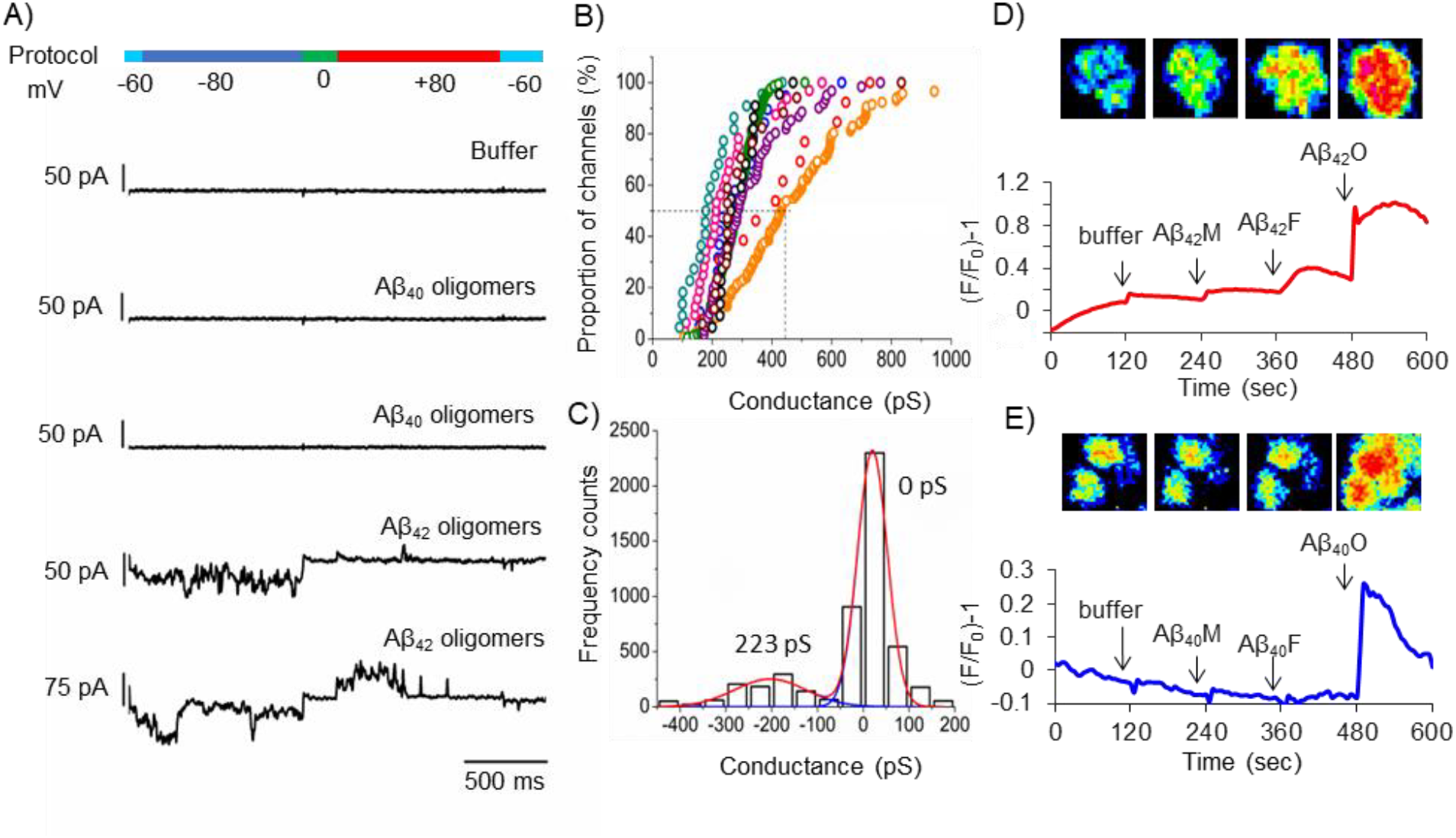
Aβ oligomer induced ion cellular influx. Patch-clamp conductance recording, voltageclamped, for membrane excised from HEK293 cells. Only Aβ_42_ oligomers (lag-phase) but not Aβ_40_ shows strong conductance (A). Conductance form nine individual excised membrane, each data point is the modal conductance (pS) with channel from a 1000 ms fixed voltage (B). Distribution of conductance’s over a single 1000 ms interval (C). Detection of intra-cellular Ca^2+^ in HEK293 cells by total Fluro4 florescence. Also shown, single-cell florescence imaging of HEK293 cells after addition of buffer; monomer (M); fibril (F); oligomer (O) (left to right) Aβ_42_ (D) and Aβ_40_ (E). Only oligomers of both Aβ_42_ and Aβ_40_ show Ca^2+^ influx detected by Fluro4 florescence. Box-size for cell images; 20 µm and 30 µm, for panel D and E respectively.

We were particularly interested in the magnitude of the conductance because this can be related to the dimensions of the channel, using the Hille equation(53, 54), see methods. The conductance values observed during the patchclamp recordings, have been ranked in size for each individual excised membrane. Conductance’s for nine individual excised membrane measurements are shown in Figure 6B. Each data point in Figure 6B is derived from a 2500 ms period of measurement, which are analysed as conductance distribution plots, an example of which is shown in Figure 6C. For the nine-patches analysed in detail, Table S1 shows the median and upper and lower limit of conductance values. The median value for the nine channels is 320 pS. There is also considerable variation in the values for each channel, a typical range of conductance, observed between 80 % and 20 % of the range of values, is for example, 320 +/-170 pS. Some patch-clamp measurements hint at multiple channels with distinct pore sizes, Figure 6B.

Assuming a channel length of 54 Å (Figure 5) which is sufficient to span the lipid membrane, then a channel conductance of 320 pS, suggests a channel diameter of 14 Å, which is similar to that indicated in the cryo-EM structure, Figure 5. The range of conductances typically observed for a single channel is between 200-500 pS, which suggests a channel diameter of 11-19 Å respectively (see Table S2). The rapid changes in conductance, observed in the patch clamp measurements, suggests the channel is capable of rapid movement in diameter. The similarity in predicted and observed diameters implies a connection between the observed conductances and the Aβ_42_ annular structure.

Aβ oligomers and curvilinear protofibrils can also cause much smaller conductances, described as a general membrane permeability, which has previously been reported across a synthetic monolayer(43). These are not apparent in the patch-clamp recordings of excised membranes shown here. However, these can be detected by Ca^2+^ influx measurements. We have used, Fluro-4, an intracellular Ca^2+^ sensitive fluorescent dye, to measure Ca^2+^ influx, Figure 6D, E. Unlike the Aβ ion-channel conductances, the permeability to Ca^2+^ happens almost instantly after exposure from Aβ oligomers. The spike in intracellular Ca^2+^ reaches a maximum within 60 sec, the Ca^2+^ signal is then partially lost as the fluorescent dye leaks from the cell, due to the presence of Aβ oligomers, shown in Figure 6D, E and supplemental Figure S10. While addition of Aβ_42_ monomers and fibrils have minimal measurable impact on levels of Ca^2+^ influx into the cellular lumen, Figure 6D, E. Furthermore, Ca^2+^ influx is caused by both Aβ_42_ and Aβ_40_ oligomers, while the large (200-500 pS) ion channels are only observed for Aβ_42_ in cellular membranes and not Aβ_40_, Figure 6A.

Cryo-EM and TEM images of lipid vesicles indicate prefibrillar Aβ oligomers and curvilinear protofibrils embed and disrupt the lipid bilayer within 30 seconds after exposure to Aβ, Figure S11 and S12. The effect of the annular assemblies is very different; it typically takes 10 minutes for channels to form in the membrane and the channel might remain active and open for half an hour or more. For these reasons we suggest this spike of Ca^2+^ influx is not the same phenomena as the ion conductance measured by patch-clamp. This implies that the instant Ca^2+^ influx is not caused by annular structures, but is the result of curvilinear protofibrils inserting and carpeting the membrane, which we have previously imaged(13) and now show in Figure S11 and S12 with an incubation time of just 30 seconds.

## CONCLUSION

Our cryo-EM and cryo-ET imaging of the prefibrillar assemblies has enabled characterization of this group of key structures in near native aqueous conditions. The collection of structures observed, suggests a mechanism of assembly, which points to an extension from small oligomers to curvilinear protofibrils, together with lateral association of these to form fibrils. Furthermore, the dimensions of the annular structures, imaged in an aqueous environment, makes a strong link to ion-channel conductance measurements. In the future, we hope it may be possible to obtain atomic resolution structures, imbedded in lipid-bilayers.

More generally, annular structures are formed for as many as six different amyloid-forming proteins(40). Annular structures, believed to be responsible for Parkinson’s Disease, have been reported for α-synuclein and imaged by cryo-EM(45). The small size and inherent flexibility of these annular assemblies has meant that only low-resolution structures have been determined. Like Aβ, α-synuclein has also been shown to form ion channel pores(55, 56). Molecules which block the annular channels or inhibit their formation are potential therapeutics for treating AD(38, 57, 58).

## Materials and Methods

### Aβ sample preparation

Aβ wildtype peptides (Aβ_42_ and Aβ_40_) were purchased from GenScript and EZBiolab, which were synthesized by F-moc ((*N*-(9-fluorenyl) methoxycarbonyl) chemistry. The lyophilized Aβ peptides were solubilised at 0.7 mg ml^-1^ (or 2 mg ml^-1^ for cryo-EM) in ultrapure water at pH 10. Aβ monomers were isolated by size exclusion chromatography (SEC) using a Superdex 75 10/300GL column. Concentrations of Aβ stock solutions were determined by absorbance using a spectrophotometer (Hitachi U-3010) at 280 nm (ε = 1280 M^-1^.cm^-1^).

Aβ stock solution was diluted into 10 μM by buffer (160 mM NaCl and 50 mM HEPES at pH 7.4), aliquoted into a 96-well plate and then placed into a fluorescent well-plate reader (FLUOstar Omega). Well plates were set to shake for 30 seconds every 30 minutes followed by a fluorescence reading, fluorescence excitation at 440 nm and emission at 490 nm. Aβ fibril formation was monitored by thioflavin-T (ThT) fluorescent dye in separate wells. Aβ oligomers were obtained from the well plate at the end of the lag-phase, with ThT in adjacent wells. The lag-phase oligomer preparations were immediately used or stored at -80 °C for use in cryoET, patch-clamp measurements and cellular Ca^2+^ influx measurements.

#### Negatively stained screening with Transition Electron Microscopy

The Aβ preparations were characterised by negatively stained electron microscopy. Aβ_42_ oligomers (10-30 µM monomer equivalent) were loaded onto glow-discharged carbon-coated grids with a 300 mesh, by a droplet (5 μl) method and washed with ultra-purified water. Followed by uranyl acetate staining (2% (w/v), pH 7.4) for 10 sec, before a final wash step. Images were taken with a JEOL JEM-1230 electron microscope operated at 80 kV, using the Olympus iTEM software.

### Cryo-Electron Microscopy (Cryo-EM)

#### Aβ_42_ oligomer preparation for cryo-EM

Lyophilized peptides were solubilised at 2 mg ml^-1^ in ultrapure water at pH 10. Aβ42 monomer (300 μM) were incubated with NaCl (160 mM) and HEPES (50 mM) buffer at pH 7.4 at room temperature, after vertexing for 15 secs. Aβ_42_ prefibrillar assemblies with predominantly oligomeric and curvilinear protofibril structures, were obtained after 15 mins incubation, confirmed by negatively stained electron microscopy. ‘Lag-phase’ oligomeric samples were ready to be loaded onto cryo-grids immediately.

#### Cryo-EM

Holey carbon grids (Quantifoil Au R 2/2, 200 mesh) were glow-discharged with easiGlow at 20 mA for 45 s. Aliquots (4 μl) of Aβ_42_ oligomers were applied to the grids and blotted for 2.6 s with filter paper (Whatman) at 75% humidity and 4°C using Leica EM GP2 plunge freezer. Sample grids were prescreened on Jeol JEM-2100Plus. The dataset was acquired on a Thermo Fisher Titan Krios microscope, 300 kV, with Gatan K3 detectors. Data was acquired with an aberration-free image-shift, using EPU software. In total, 13,843 movies were collected, at super resolution 0.55 Å/pixel, with a total dose of 40 e/Å^2^ and defocus range between 0.5 to 3.5 μm.

#### Data Processing

All super-resolution frames were gain-corrected, binned, aligned, and dose weighted, this was followed by summation into a single micrograph using motion-correction in RELION. The pixel size was binned to 1.1 Å/pixel. The estimation of parameters for the contrast transfer function (CTF) was carried out using CTFFIND4.1 in RELION. Initially, manually selected particles from 50 micrographs were used to generate a model for automatic particle picking using Cryolo. Further processing was performed in CryoSPARC. Particles were binned and extracted at 2.2 Å/pixel. 2D classification was performed from 3 million extracted particles. A workflow is shown in supplementary Figure S13. *Ab-initio* 3D models were reconstructed *de novo* from 2D class averages. Non-uniform refinement was used to improve the resolution. Molecular graphics and analyses were performed in Chimera (UCSF). A protofibril model was fitted into an S-shaped half-map of a mature fibril, pdb: 2MXU(19).

### Cryo-electron tomography (Cryo-ET)

Lag-phase oligomers of Aβ_42_ (5 µM, monomer equivalent) in HEPES buffer (30 mM) NaCl (160 mM), were applied to Quantifoil R2/2 holey carbon grids and then plunge-frozen using a Thermo Fisher Vitrobot. Cryo-electron tomography (cryo-ET) images were obtained with a Thermo Fisher Glacios TEM operating at 200 kV, using a 4k x 4k Falcon 3EC direct electron detection camera, with a magnification of 73k. Pixel size was 1.9 Å at the specimen level. Tilt series were obtained between −60° to +60° with increments of 3° using the dose symmetric scheme. The defocus was set between 3 and 4 µm, and the total dose was *ca*. 100 e/Å^2^. Typically, tomographic images shown are a sum of 10 slices, 7.6 nm thick. 3D tomograms were reconstructed from the tilt series using RAPTOR(59) and the IMOD software package, followed by Weighted Back Projection (WBP) or Simultaneous Iterative Reconstruction Technique (SIRT), with the TOMO3D package(60, 61). Final tomograms were binned 4x, resulting in a pixel size of 7.6 Å. Images of a single threshold surface were produced using Chimera (62) following a 3×3×3 median filtering operation. The lengths of curvilinear protofibrils, in three dimensions, were obtained by manual inspection in Fiji(63) of volume regions previously rotated and extracted from full tomograms in 3dmod (slicer module).

### Patch-clamp conductance measurements

#### Cell culture

HEK293 immortal cells were incubated at 37 °C, 5% CO_2_ incubator, in Dulbecco’s Modified Eagle Medium (DMEM) supplemented with 10% fetal bovine serum (200 units) and penicillin-streptomycin (0.2 mg/ml) in a 30 ml cell culture flask. Cells were split at ∼70-80% confluency around every 5 to 7 days using Ca^2+^ and Mg^2+-^free phosphate-buffered saline (pH7.2) during which, a fraction of cells were plated in 35 mm diameter easy-grip Petri dishes with same buffer and incubated for 2-3 days before use. Reagents and media were purchased from Sigma-Aldrich and ThermoFisher scientific (Invitrogen).

#### Patch clamp Recordings

All patch clamp measurements were made from excised membrane in an inside-out configuration. This allows Aβ to diffuse to the extracellular surface of the HEK293 cell plasma membrane, in voltage-clamp mode. Tip burnt polished micro glass pipettes (GC150TF-10, Harvard Apparatus) were backfilled with extracellular buffer (containing (mM): 121.4 NaCl, 10 CsCl, 9 NaHEPEs, 1.85 CaCl_2_, 1.87 MgCl_2_, 2.16 KCl, and 0.1 EGTA) with addition of Aβ oligomers 5 μM (monomer equivalent) at pH 7.4. Aβ oligomers were diffused towards the extracellular membrane surface within the pipette. Membrane patches of approximately 1-3 μm diameter. The pipette resistance varied from 2-6 mΩ when filled with recording solution. Junction potentials appeared at boundaries of ionic asymmetry and were considered for using an applied pipette offset potential. Recordings were sampled at a rate of 2 kHz with 500 µs intervals with a low-pass 8-pole Bessel filter frequency of 0.2 kHz. The holding potential was set to -60 mV.

Channel-induced transmembrane currents were recorded by voltage clamping for 30-45 minutes. A protocol from -80 to 0 to +80 mV using Axopatch 200B amplifier (Axon Instruments) was applied using pCLAMP 11 software. All recording was processed using Clampfit software and a lowpass boxcar filter was applied.

#### Determining Conductance values

Current/voltage was used to calculate channel conductance (pS). Distribution plots of conductance for a 2500 ms interval were plotted, see Figure 6C for an example. These were used to obtain typical (modal) conductance values of a 2500 ms intervals, and used as a single data point for the total recording, Typical 40-100 data point for each excised membrane were obtained, Figure 6B.

#### Calculation of channel pore diameter using channel conductance

To calculate the theoretical pore diameter, a model developed by Hille (53) and adapted by Cruickshank et al (54) was adopted. Pore diameter (d) was calculated by ionic conductance (g), pore length (l) and solution resistivity (ρ) of 80 Ω cm(64) (Equation 1).

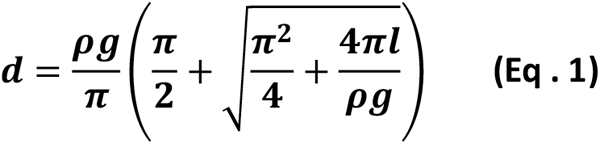

### Cellular Ca(II) influx

#### Fluo-4AM loaded HEK293 cells

HEK293 cells were maintained at 37 °C in 5% CO_2_ containing Dulbecco’s modified Eagle’s medium (DMEM, purchased from Thermofisher) supplemented with 10% fetal bovine serum and penicillinstreptomycin (0.2 mg/ml). Cells were plated into a 12-well plate, 1 mL each well, and incubated overnight, cells typically gained *ca*. 50% confluence. Next DMEM was replaced with fresh DMEM which was supplemented with 5 μM of fluo-4AM (Abcam). Plates were then left in an incubator for a further 40 min to enable the cellular uptake of the Ca^2+^ sensitive fluorescent dye. Excess extra-cellular fluo-4AM was removed by two washes of 400 μL of DMEM Eagles cell medium in each well. Cells were then incubated for a further 20 minutes at 37°C to allow de-esterification of intracellular fluo-4 to occur, which activates Ca^2+^ dependent fluorescence. Finally, the DMEM was replaced with an aqueous buffer containing (mM) 121 NaCl; 10 CsCl; 9 HEPES; 1.85 CaCl_2_; 1.87 MgCl_2_; 2.16 KCl, buffered to pH 7.4, to mimic conditions used for patch clamp recordings. The fluo-4 loaded cells were then ready for time-lapse fluorescence microscopy imaging.

#### *Ca*^*2+*^ *Fluorescence Imaging of Fluo-4AM*

Fluorescence microscopy was performed using an inverted Leica DM IL microscope with 10x or 20x magnification. The bandpass filter allowed excitation at 470 nm and emission was recorded at 520 nm. Time-lapse fluorescence images and bright-field visible light images were acquired using a charge-coupled device (CCD) camera with a temporal resolution of one image every 5 s, recordings were for 10-30 mins.

Three Aβ_42_ and Aβ_40_ preparations were studied: monomers (SEC purified); oligomers (from the end of the lag-phase); fibrils (from the plateau-phase). Stock solutions of 30 µM Aβ was added to HEK293 cells within 450 µL of buffer to produce a final Aβ concentration of 5 μM. The microscope was operated using ProgRes Capture-Pro 2.8.8 software and fluorescence intensity was measured via Image-J. Fluorescence intensities were measured by analysing the overall field, using the time series analyser V3 plug-in. Individual single cell fluorescence intensities reported similar Ca^2+^ influx behaviour. Changes in Fluo-4 fluorescence signals with time were presented as (F/F_0_)-1, where (F) is the observed fluorescence and (F_0_) is the background fluorescence at a time point just before the addition of Aβ. Typically, each condition was recorded using four separate wells, with at least three independent repeats. Ionomycin (Merck) disrupts cellular membrane integrity and was used as a positive control, to confirm consistent cellular incorporation of Fluo-4AM.

### Lipid Vesicle Preparation and Imaging

Large unilamellar vesicles (LUVs) were produced using an extrusion method. The lipids used a mixture of 68:30:2 by weight of phosphatidylcholine (PC): cholesterol: monosialotetrahexosyl-ganglioside (GM1), for further details see (13). After air drying, the lipid film was resolubilized in aqueous buffer, NaCl (160 mM) HEPES (30 mM) buffered at pH 7.4, to a lipid concentration of 1 mg mL^-1^. LUVs were formed from solubilized lipids using a benchtop mini extruder (Avanti Polar Lipids, Alabama, USA). The lipid solution was passed across a polycarbonate membrane (Cytiva Whatman, USA), with a 100 nm pore size, 21 times at a temperature of 65 °C.

Cryo-EM imaging of the vesicles were in the absence and presence of Aβ42 oligomers (20 μM monomer-equivalent, obtained at the end of the lag-phase) incubated with the lipid vesi-ccles (0.5 mg mL^-1^) for just 30 seconds. The lacey carbon grids (300 mesh) (Quantifoil) were glow-discharged with easiGlow at 20 mA for 45 seconds. A 4 μL aliquot of the vesicles were applied to the grids and blotted for 2.4 second with filter paper (Whatman-level 1) at 75% humidity and 4 °C using a Leica EM GP2 plunge freezer. JEM-2100Plus (JEOL, Ltd., Japan), operated at 200 kV, fitted with a GATAN OneView camera was used to image the vesicles.

Vesicles (0.05 mg mL^-1^) were also imaged by TEM negatively stained in the absence and presence of Aβ42 oligomers (10 μM monomer-equivalent, obtained at the end of the lag-phase) incubated with the lipid vesicles for just 30 seconds. Then 5 µl aliquots of sample were added to glow discharged Lacey carbon-coated 300-mesh grids (Agar Scientific Ltd) by the droplet method, then blotted after 30 seconds and rinsed with ddH_2_0. Following this, (4 µL) uranyl acetate (1 g/100 mL) was added, then blotted after 15 seconds and rinsed with ddH_2_O. Images were taken from a JEM-2100 plus electron microscope (JEOL, Ltd., Japan), operated at 200 kV, fitted with a GATAN One-View camera at 50,000x Magnification.

## AUTHOR INFORMATION

## Author Contributions

All authors have given approval to the final version of the manuscript.

## ACKNOWLEDGMENT

We are thankful for the support of the BBSRC; project grant code BB/Y001931/1 and BB/M023877/1. Thanks to N. B. Cronin for assistance with cryo-EM image acquisition, LonCEM Facility, The Francis Crick Institute, London, UK.

## ABBREVIATIONS

Aβ: Amyloid-beta
cryo-EM: cryo electron microscopy
cryo-ET: cryo electron tomography
HEK: human embryonic kidney cells

