## Supplemental Table S1-S2 and Figure S1-S13 for "Structural architecture of amyloid-β oligomers, curvilinear protofibrils and annular assemblies, imaged by cryo-EM and cryo-ET": Ab_oligomers_Supplimental_FIGURES_XIII_AS_SENT_March1st.pdf

**Supplementary Material**

Table S1-S2

Figure S1-S13

| | Conductance (pS) | | | |
|---|---|---|---|---|
| **Patch no.** | **20%** | **50%** | **80%** | **Difference (20% and 80%)** |
| 1 | 87 | 217 | 348 | 261 |
| 2 | 94 | 235 | 376 | 282 |
| 3 | 102 | 255 | 408 | 306 |
| 4 | 114 | 284 | 454 | 341 |
| **5** | **130** | **320** | **510** | **380** |
| 6 | 146 | 364 | 582 | 437 |
| 7 | 153 | 382 | 611 | 458 |
| 8 | 167 | 417 | 667 | 500 |
| 9 | 226 | 564 | 902 | 677 |
| **Average** | 135 | 337 | 539 | 405 |
| **SD** | 44 | 109 | 174 | 131 |

**Table S1: Conductance of A$\beta_{42}$ channels recorded from nine individual patches.** Taken from Figure 6B. The range of conductance's for each of the nine channels is shown as 20%, 50%, and 80% of the maximum conductance value. The channel with the median conductance is highlighted in bold. The mean and standard deviation are displayed at the bottom of the table for reference.

| Conductance (pS) | Implied diameter (nm) | |
|---|---|---|
| | 54 Å pore length | 70 Å pore length |
| 200 | 1.1 | 1.3 |
| 250 | 1.3 | 1.4 |
| 300 | 1.4 | 1.6 |
| 350 | 1.5 | 1.7 |
| 400 | 1.7 | 1.9 |
| 450 | 1.8 | 2.0 |
| 500 | 1.9 | 2.1 |
| 550 | 2.0 | 2.2 |
| 600 | 2.1 | 2.3 |
| 650 | 2.2 | 2.4 |
| 700 | 2.3 | 2.5 |

Table S2 **Predicted Inner Diameter of channel calculated from Conductance.** The implied diameter (nm) has been calculated for conductance's between 200 and 700 pS, with two possible channel pore lengths of 54 and 70 Å.

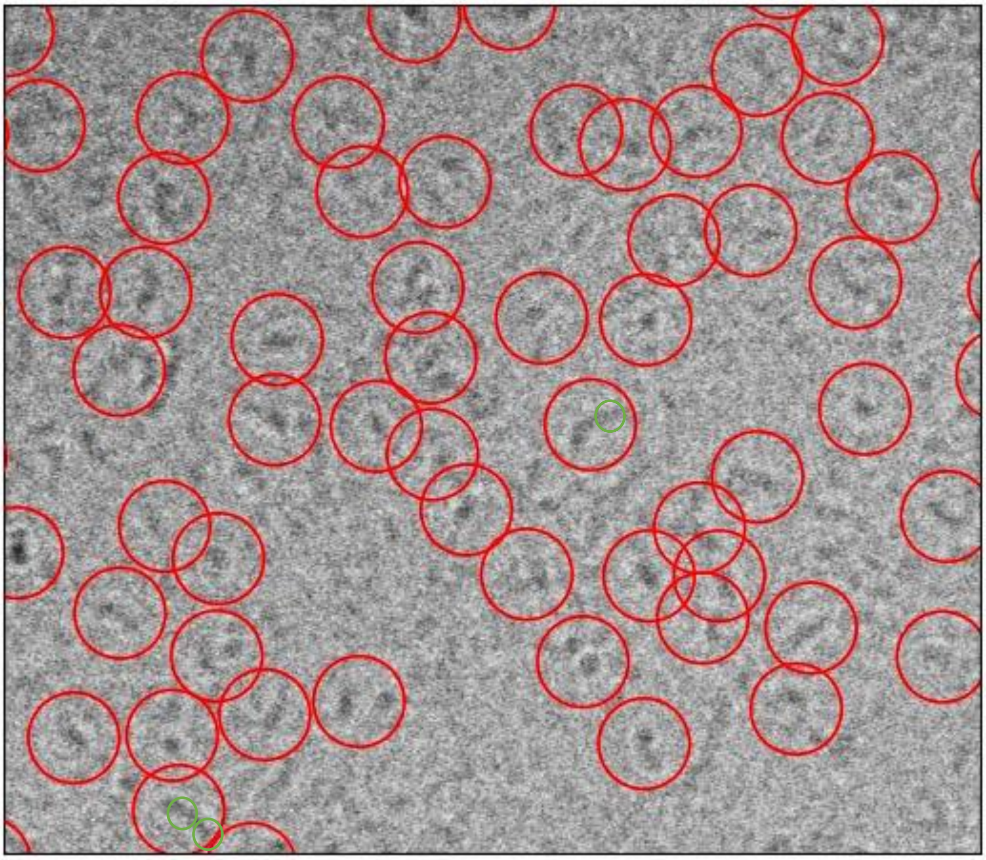

**Supplemental Figure S1**: Typical cryo-EM micrograph, circles are set at 100 Ȧ, highlighting the range of oligomers and curvilinear protofibrils. Three million particles were auto-picked, all the various forms of Aβ particles were picked simultaneously using Cryolo. (Template auto-picking was not used). Lag-phase Aβ42 assemblies 300 µM incubated for 30 mins.

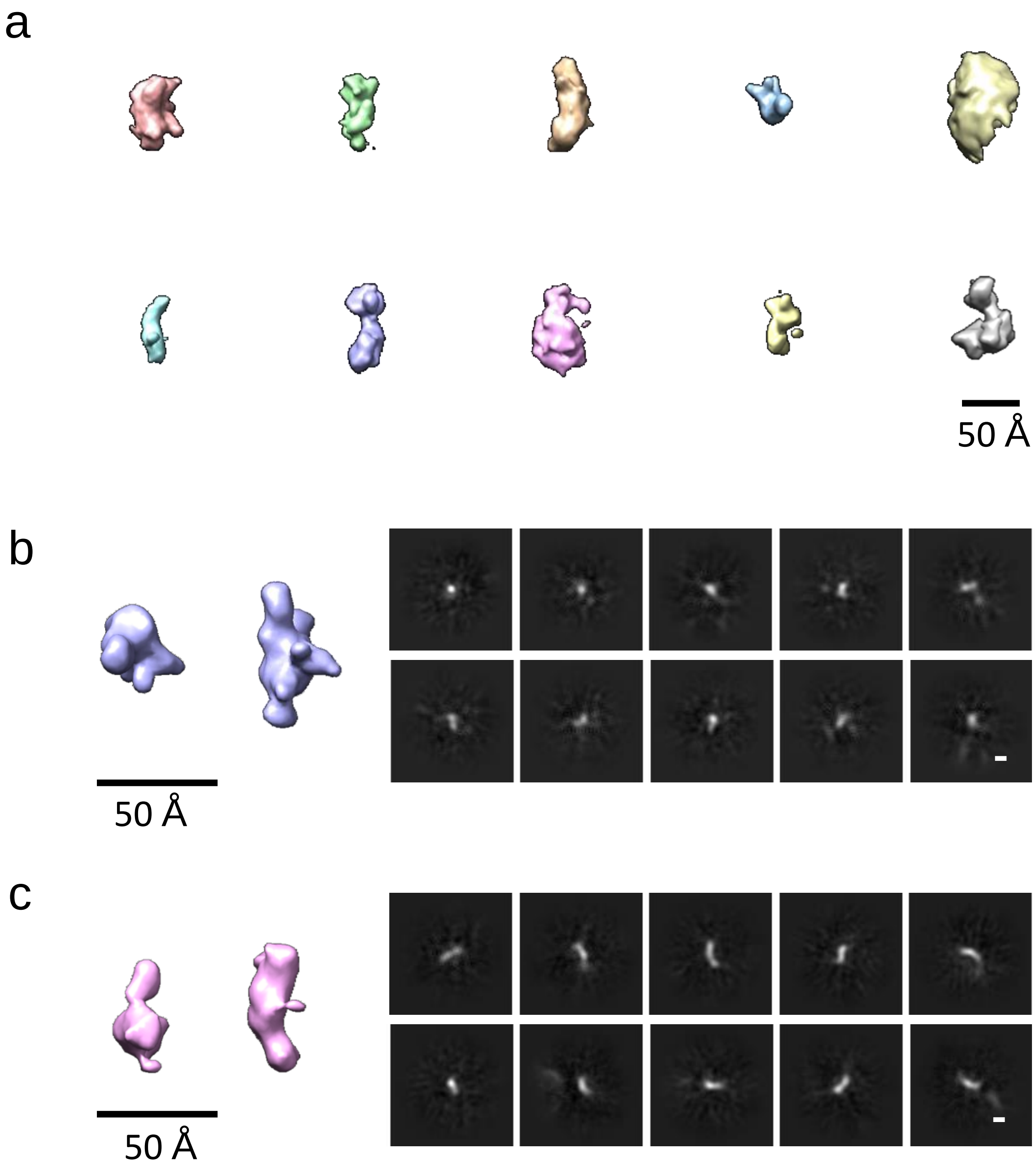


**Supplemental Figure S2a-c:** 3D *ab-initio* reconstruction for A$\beta_{42}$ oligomers, prefibrillar assemblies. a) Ten unique 3D structures generated from 2.5 million particles; between 176k and 499k particles for each structure b) & c) two individual 3D *ab-initio* models representing the smaller oligomer/protofibrils, together with representative 2D averages. Scale bar: 50 Å for 3D structures, 20 Å for 2D averages.

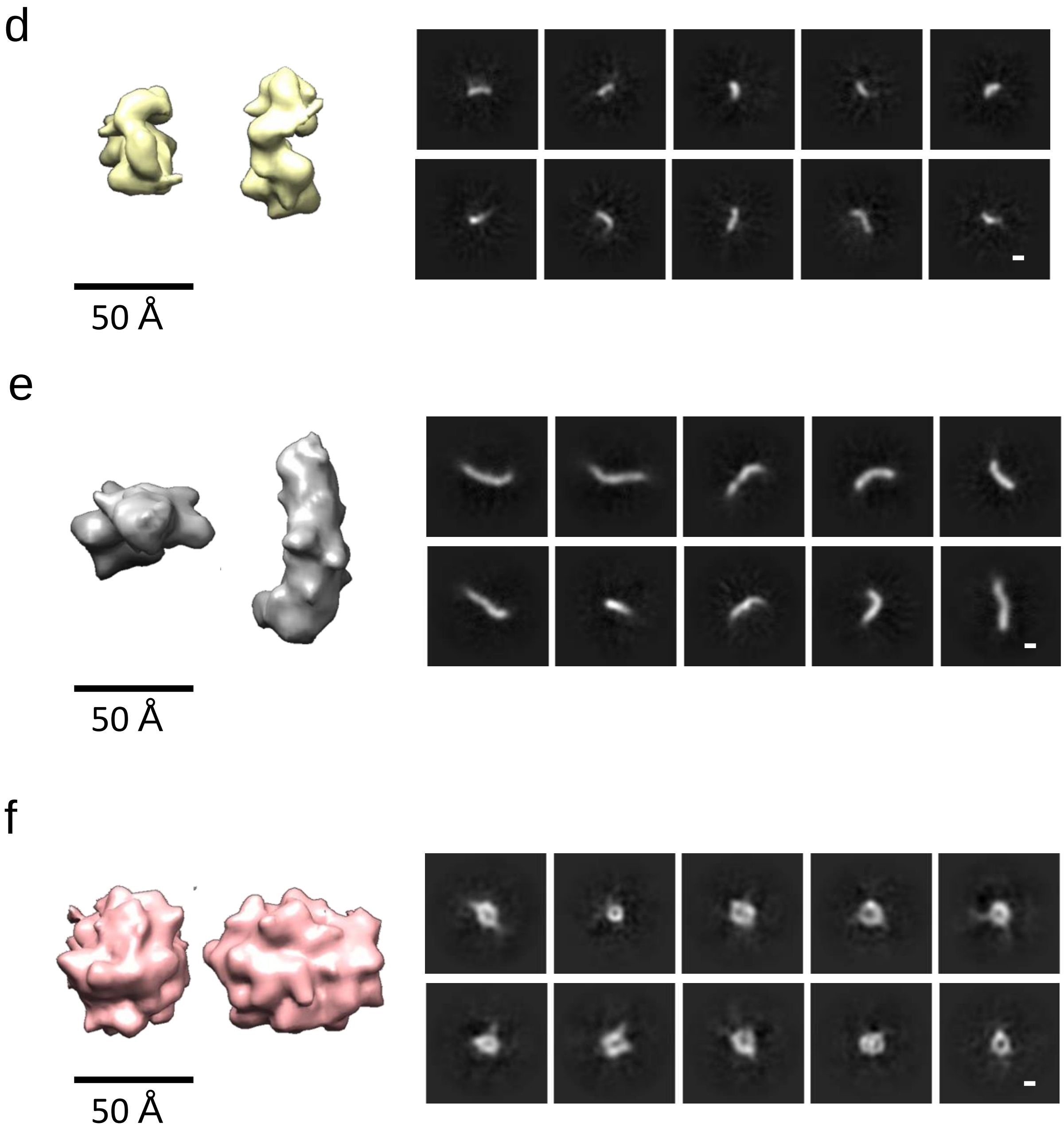


**Supplemental Figure S2d-f:** 3D *ab-initio* reconstruction for Aβ42 oligomers, prefibrillar assemblies. Showing three of the ten structures generated. d) & e) protofibrils, together with representative 2D averages. f) larger assemblies with all dimensions greater than 50 Å, for this structure, some of the 2D averages have a ring like appearance. Scale bar: 50 Å for 3D structures, 20 Å for 2D averages.

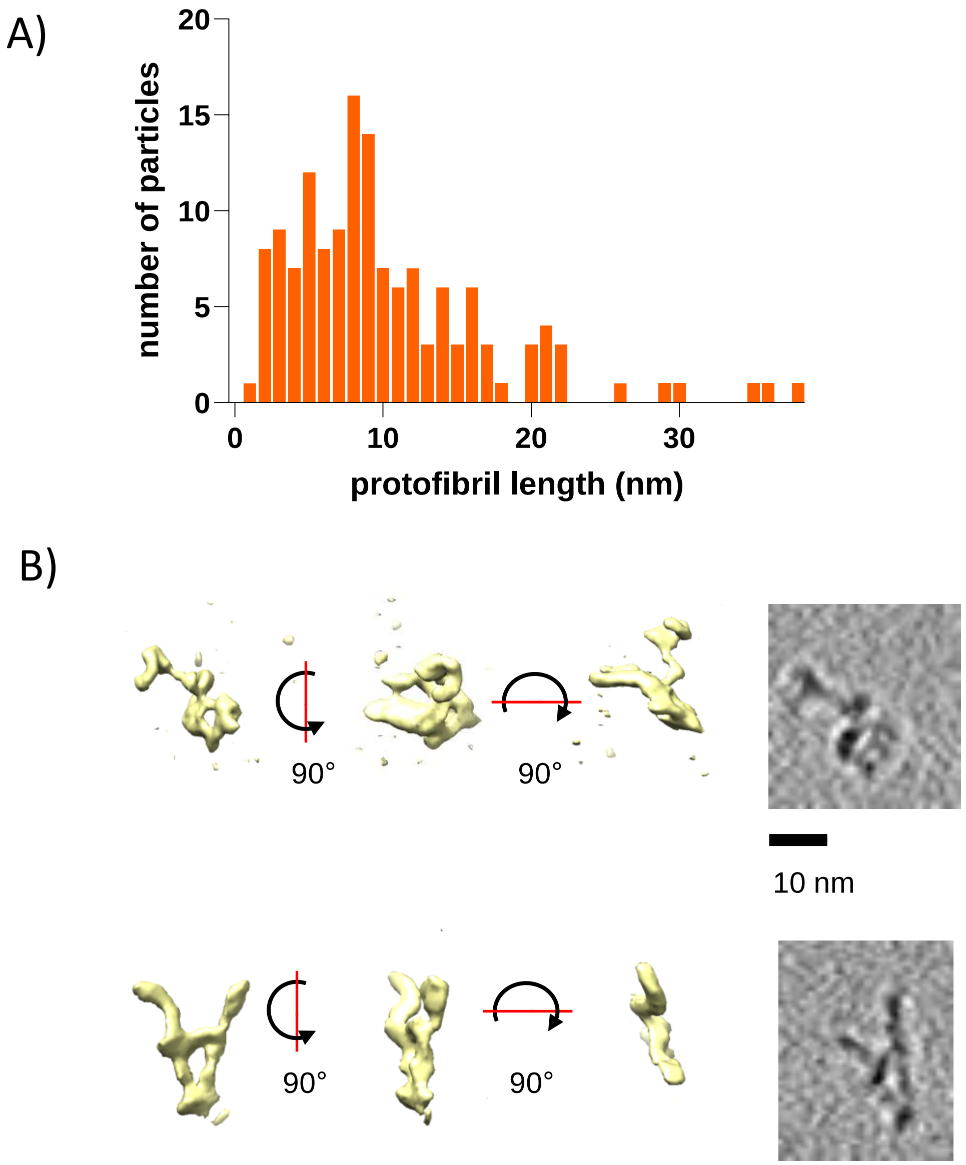


**Supplemental Figure S3:** Cryo-ET data. Distribution of length (nm) for individual protofibril particles. The majority of protofibrils are less than 9 nm long but consistently *ca*. 2.8 nm in diameter (A). Examples of branching for curvilinear protofibril assemblies, tomogram slices 7.8 Å thick, scale bar 10 nm (B).

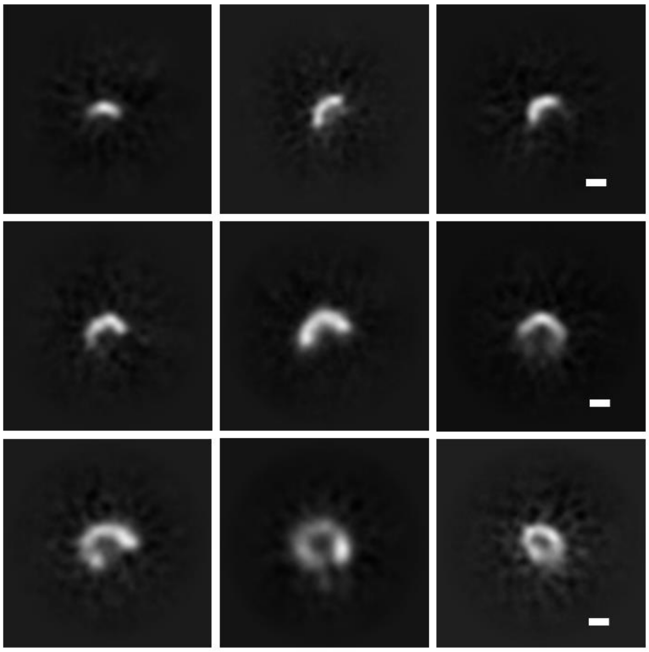

**Supplemental Figure S4** . Possible incomplete annular formation. Selected curvilinear protofibrils with a marked curvatures and ordered by length. Cryo-EM 2D class averages of *ca*. 21,000 particles per image, scale bar 20 Å.

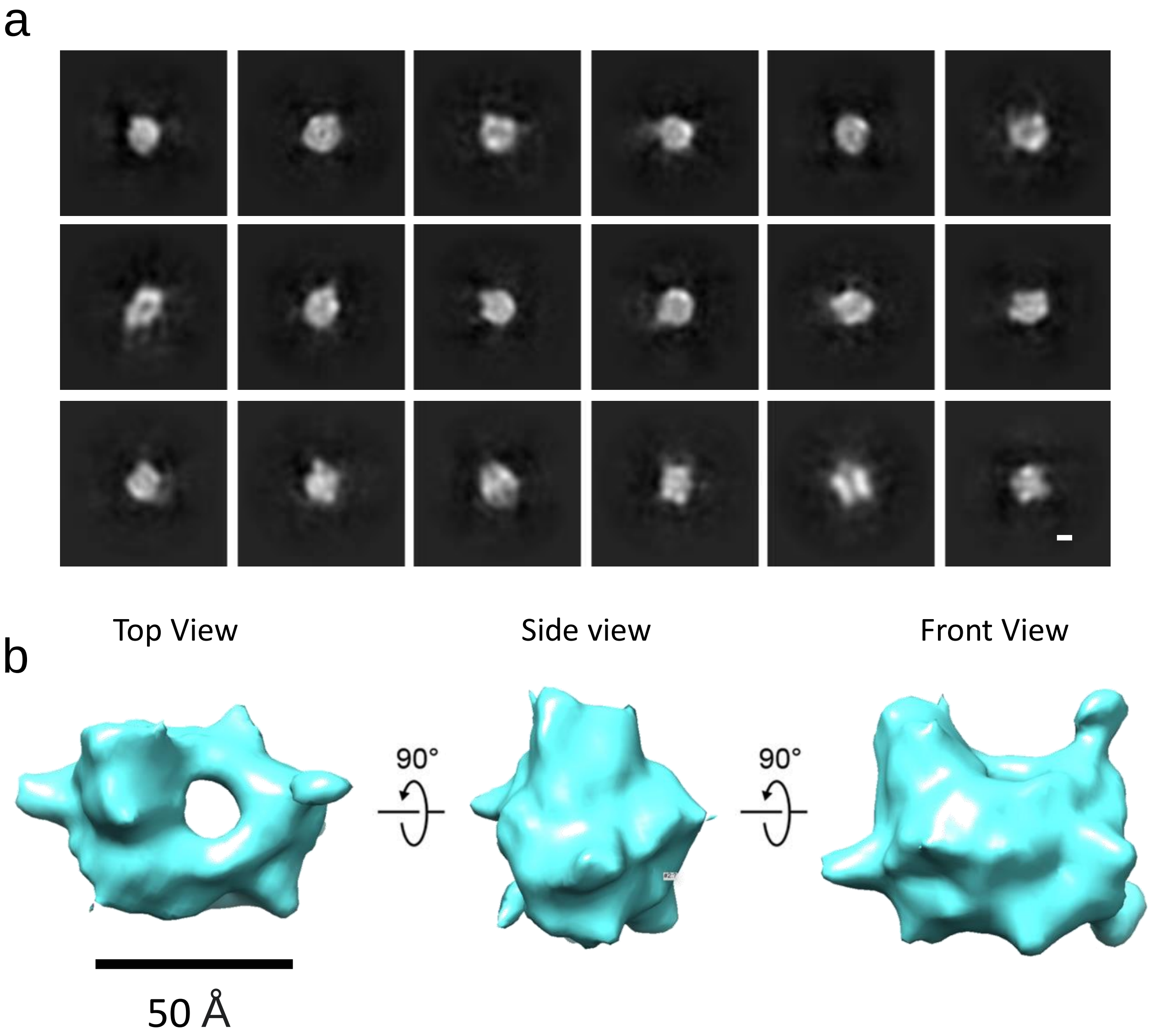


**Supplemental Figure S5.** 3D reconstruction of annular assemblies. 2D averages of 144,593 particles (a) and 3D reconstruction (b). Annular structure with an internal pore, through the middle of the structure. Dimensions of 3D: 86 Å; 74 Å; 64 Å . Scale bar: 50 Å for 3D structures, 20 Å for 2D averages.

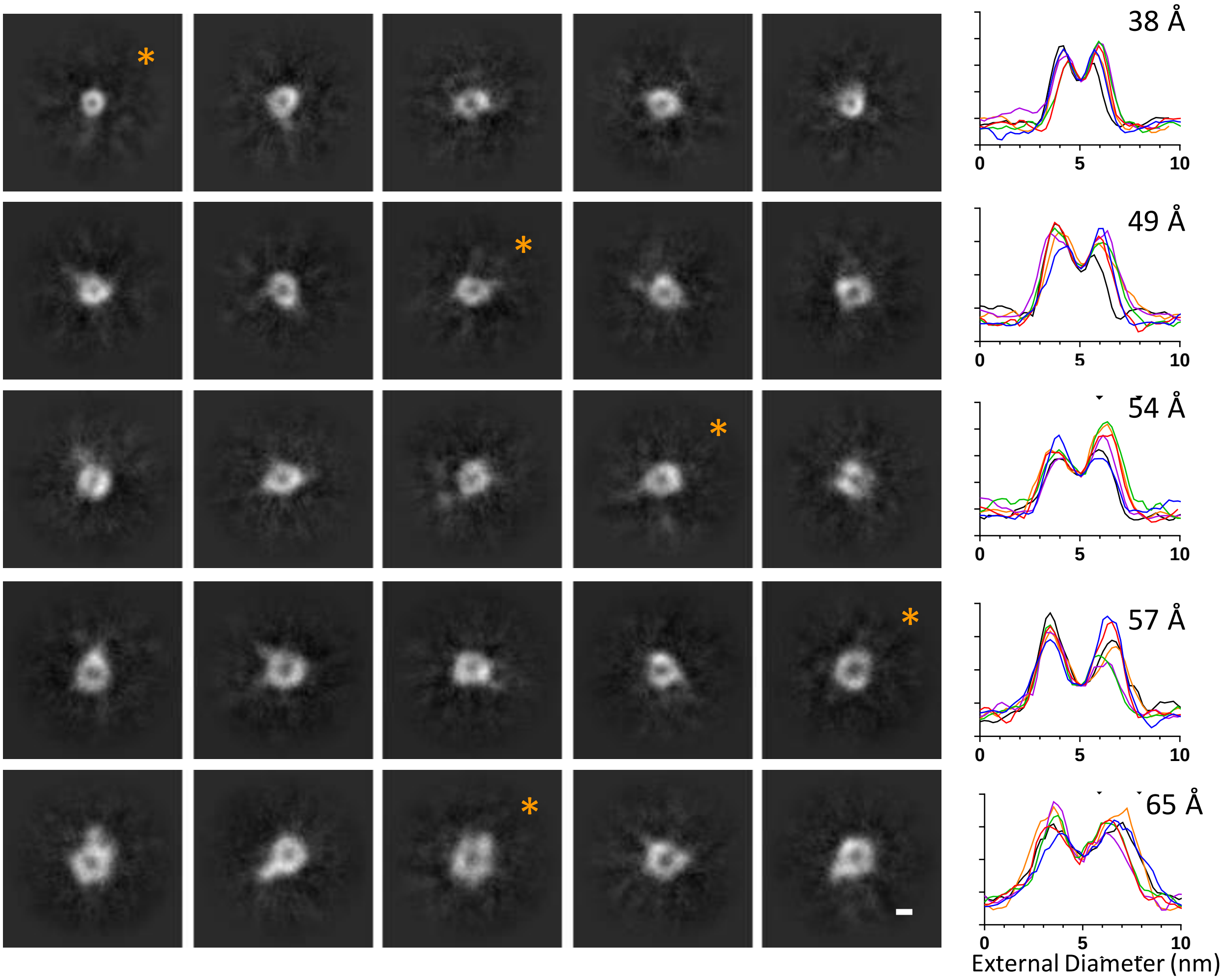


**Supplemental Figure S6. Annular oligomers top view.** 2D class averages for top (and bottom) views of annular oligomers, ranked by size, with the smaller rings on the top rows. Each class average contains ~400 particles; 10,000 particles in total. Scale bar 20 Å. To the right shows density profiles for representative annular oligomers top view, labeled by *orange, one from each row. Six profiles for angle 0, 30, 60, 90, 120, and 150 degree in colour red, orange, green, blue, black and purple. The external diameter measure from baseline of the plot profiles (row 1 to 5): 38, 49, 54, 57 and 65 Å.

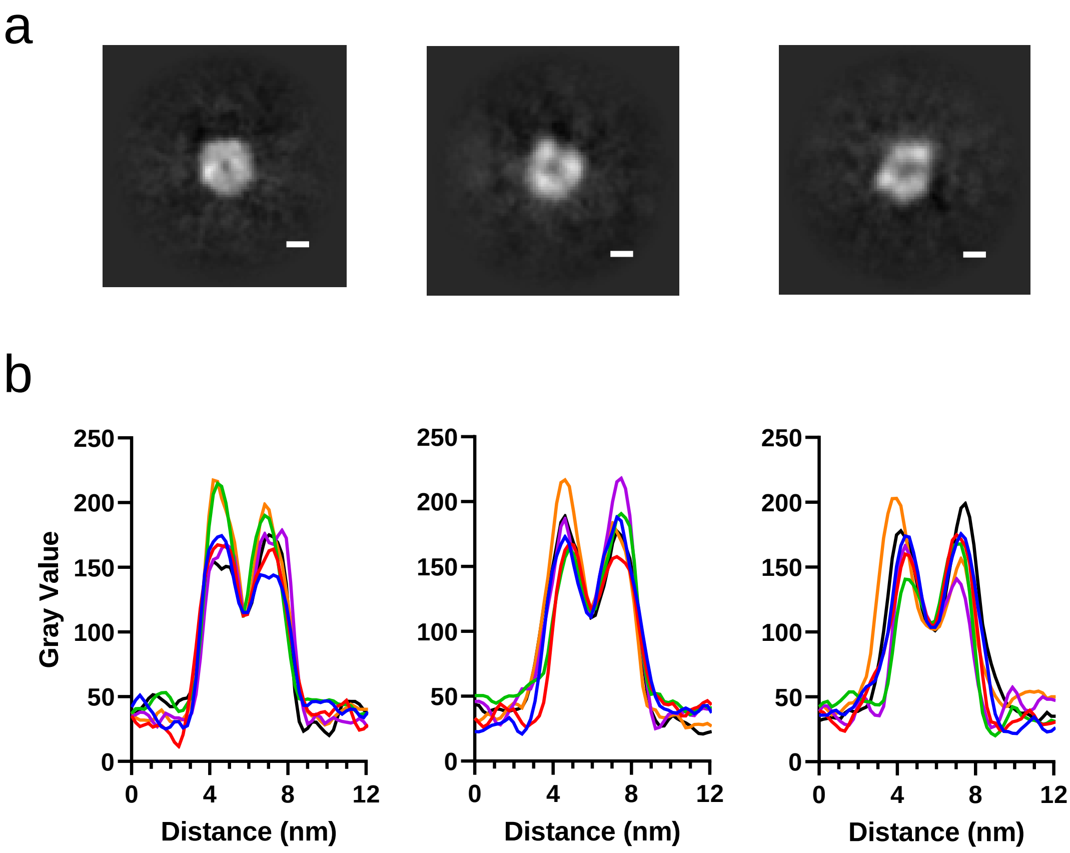


**Supplemental Figure S7.** Cryo-EM 2D class averages and density profiles of annular particles. a) The representative class averages of top view of annular oligomers. b) density profiles across the annular oligomers top view. Six profiles for angle 0, 30, 60, 90, 120, and 150 degree in colour red, orange, green, blue, black and purple. Scale bar : 20 Å.

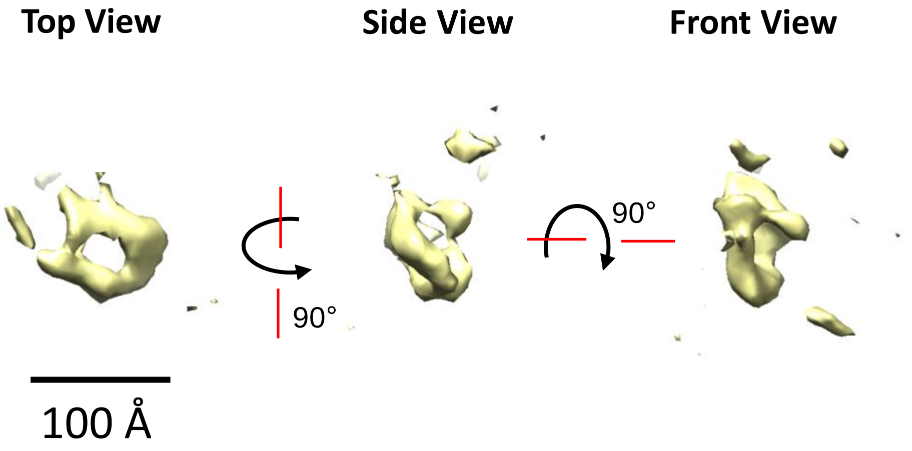


**Supplemental Figure S8:** CryoET 3D image of A$\beta_{42}$ annular assembly in near native conditions.  External diameter of ring *ca.* 70 Å, length of channel 40 Å.

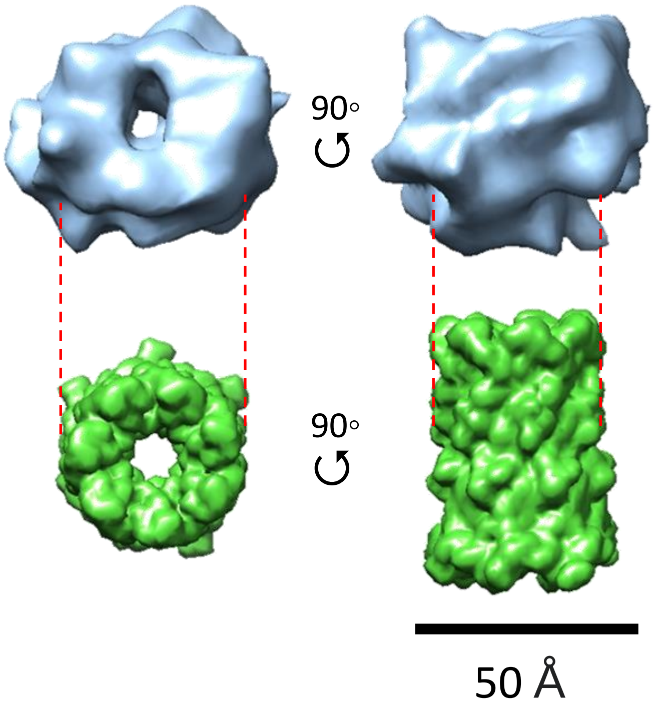


**Supplemental Figure S9.** Comparison of annular oligomer of A$\beta_{42}$ wild-type (Blue) with a Aβ42 fixed as a heptameric β-hairpin structure with α-hemolysin as a scaffold (Green) REF pdb=7O1Q. The annular oligomers of wild-type A$\beta_{42}$ is wider but not as longer as the β-hairpin contain seven A$\beta_{42}$ molecules. The density for the scaffold dependent β-hairpin contains all 42 amino acid, which may not be the case for the cryo-EM image of A$\beta_{42}$ annular oligomers. Scale bar: 50 Å.

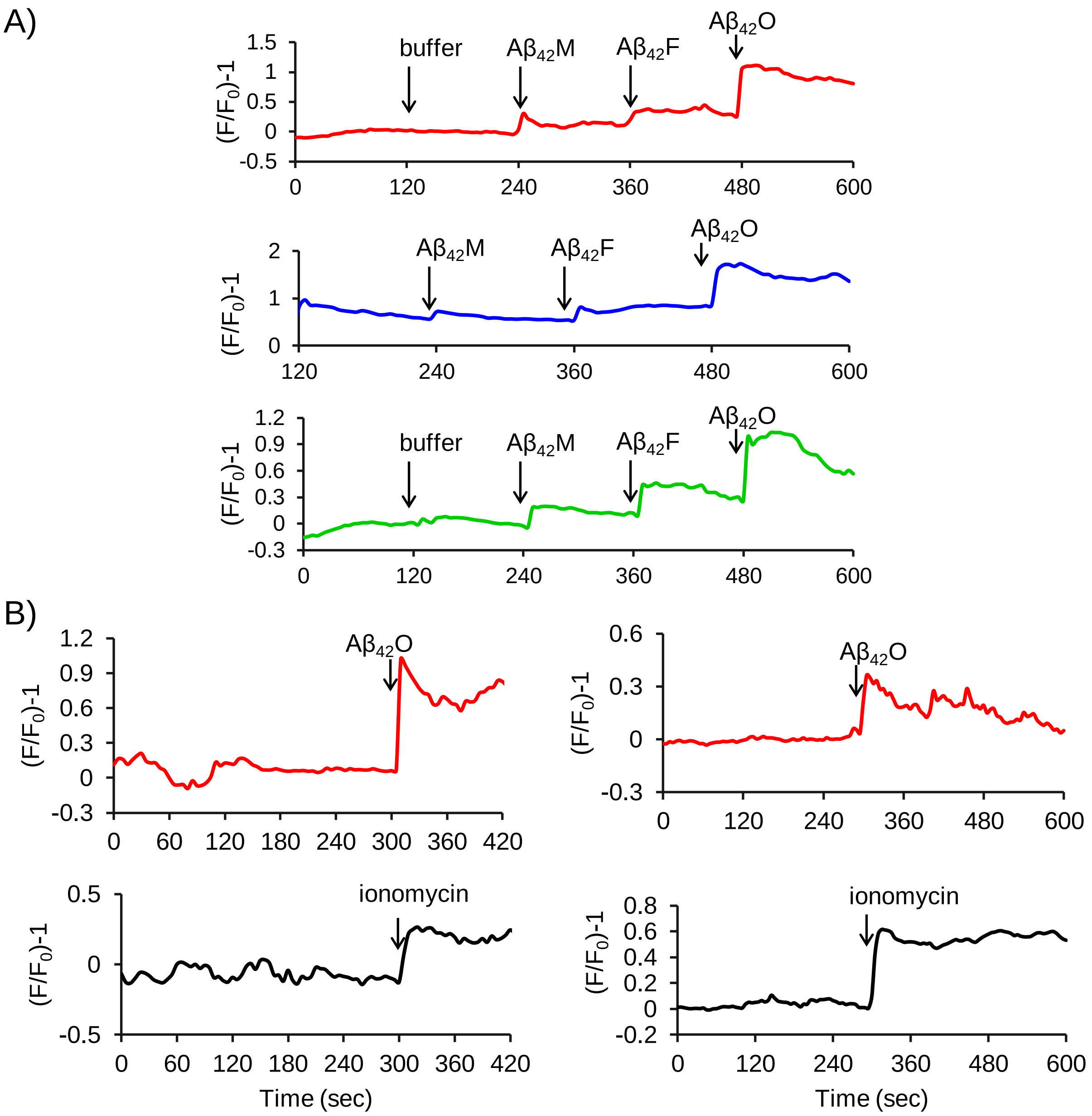


**Supplemental Figure S10:** Cellular $Ca^{2+}$ detection in HEK293 cells by Floro-4. Comparison of impact of just buffer, $A\beta_{42}$ monomer, fibril and finally oligomers (A). Comparison of impact of $A\beta_{42}$ oligomer with ionomycin. Final $A\beta_{42}$ concentration 5 μM, pH 7.4.

a) b)

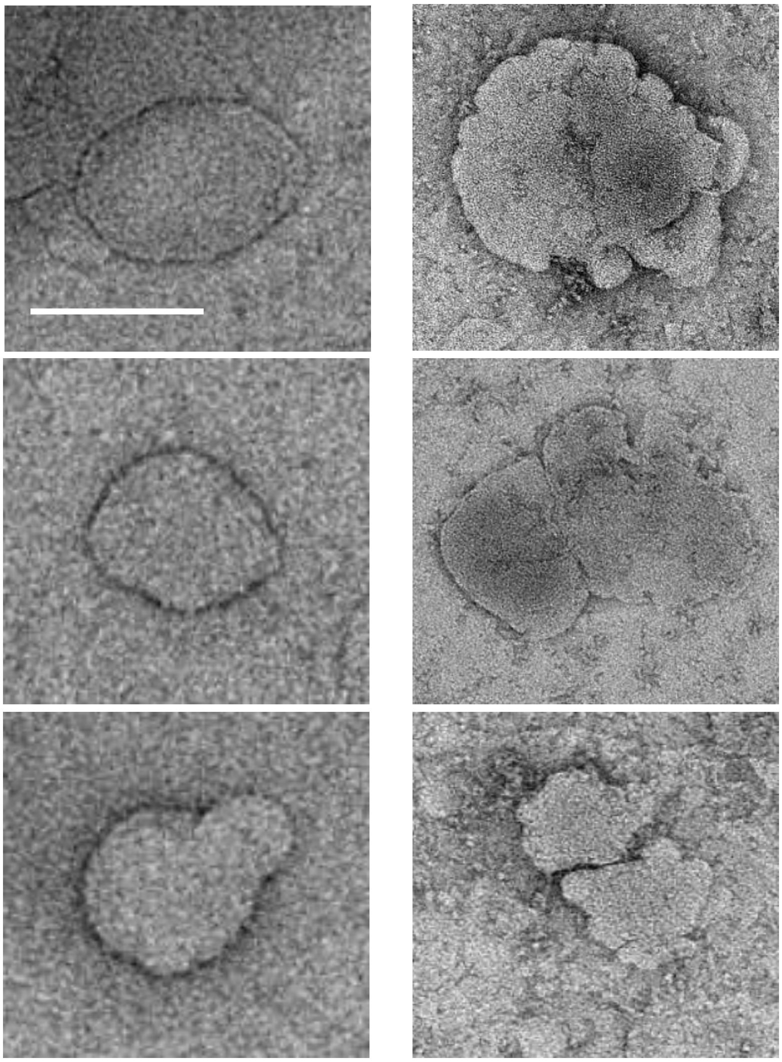

**Supplemental Figure S11:** TEM images of lipid vesicles without (a) and with the presence of A$\beta_{42}$ pre-fibrillar assemblies (b). A$\beta_{42}$ oligomers and curvilinear protofibrils incubated with LUVs for just 30 s. A$\beta_{42}$ prefibrillar assembles were obtained at the end of the lag-phase during fibril formation. Widespread distortions in the appearance of the vesicles are apparent in the presence of oligomers and curvilinear protofibrils in less than 1 min after addition of Aβ. LUVs contain PC:cholesterol:GM1 (68:30:2 by weight). Images are negatively stained with uranyl acetate. Scale bar 100 nm.

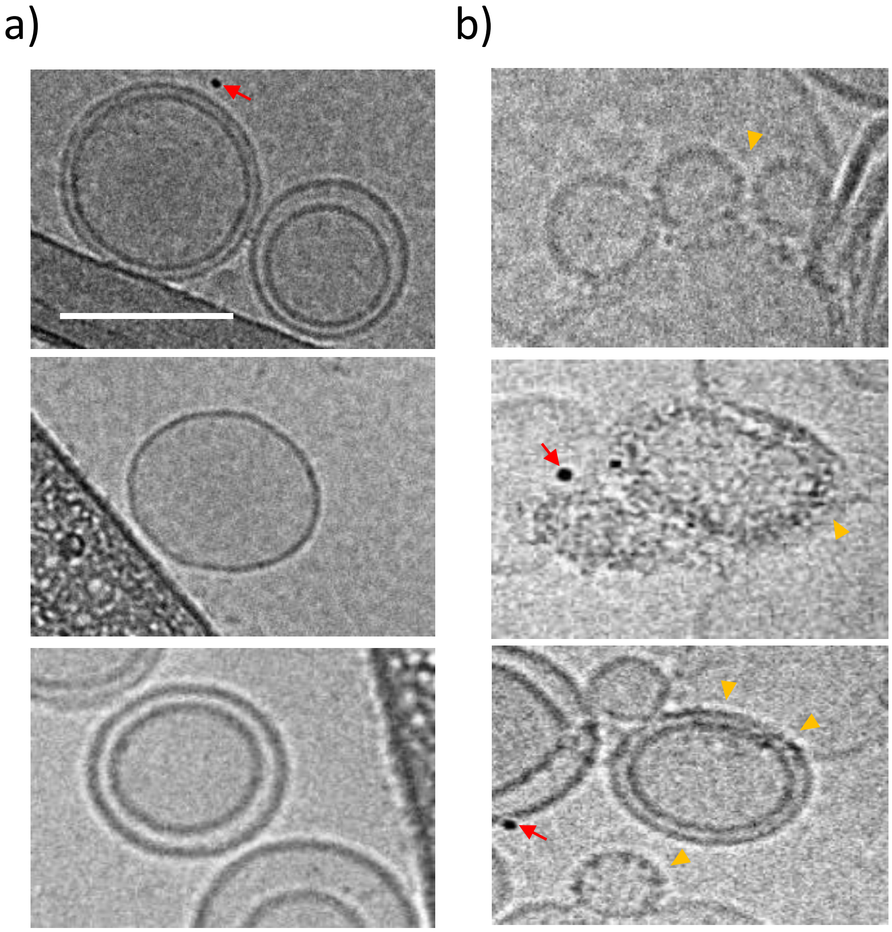


**Supplemental Figure S12:** Cryo-EM images of lipid vesicles without (a) and in the presence of A$\beta_{42}$ pre-fibrillar assemblies (b). A$\beta_{42}$ oligomers and curvilinear protofibrils incubated with LUVs for just 30 s. Within less than a minute the vesicle surface shows Aβ assemblies in the lipid bilayer (yellow triangle). LUVs (0.5mg mL$^{-1}$) contain PC:cholesterol:GM1 (68:30:2) by weight. Scale bar 100 nm, Magnification 50,000x. Red arrow represents a gold fiducial marker.

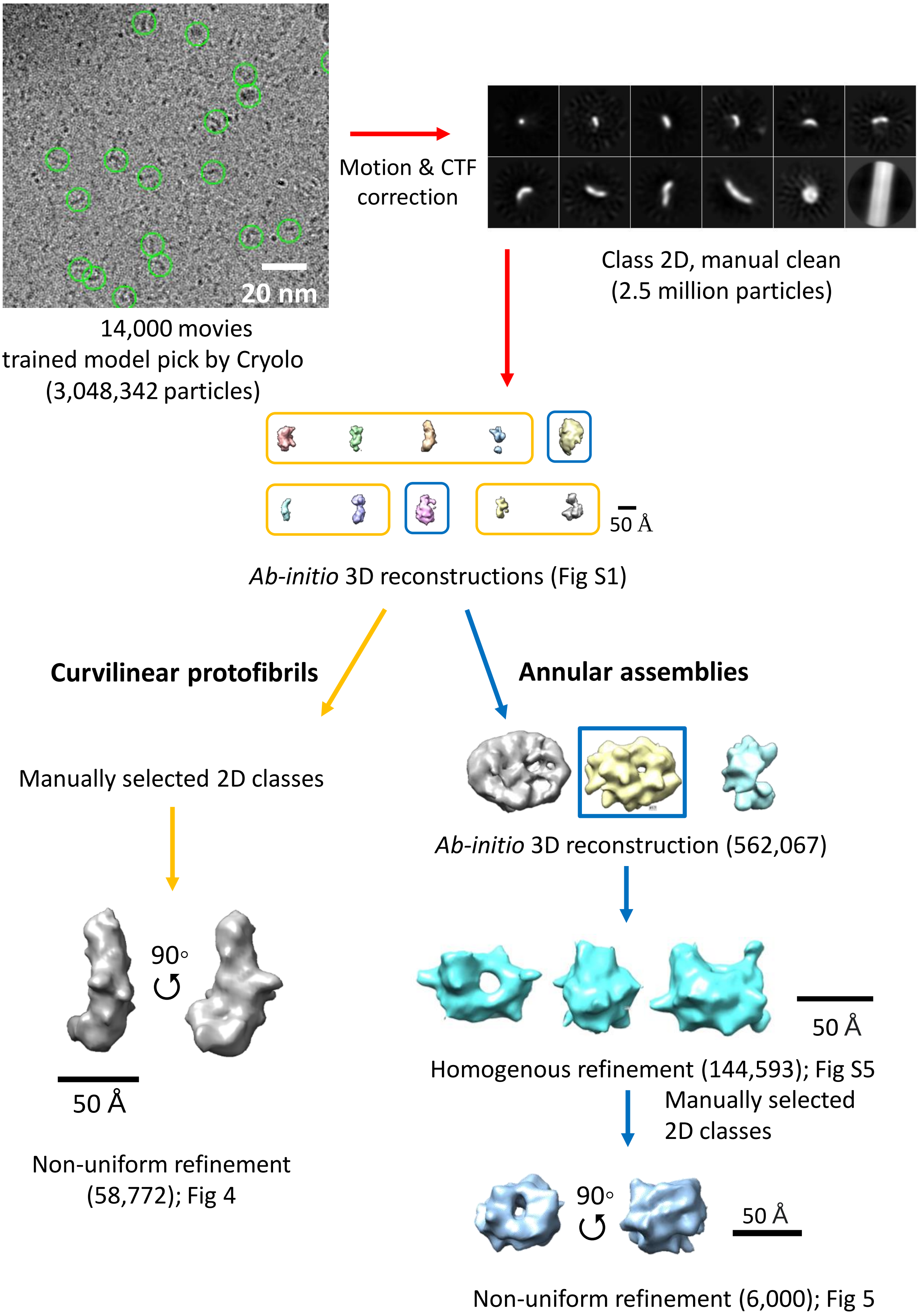


**Supplemental Figure S13:** Cryo-EM, Single Particle Analysis- Workflow
